# A new genetically tractable non-vertebrate system to study complete camera-type eye regeneration

**DOI:** 10.1101/2024.01.26.577494

**Authors:** Alice Accorsi, Brenda Pardo, Eric Ross, Timothy J. Corbin, Melainia McClain, Kyle Weaver, Kym Delventhal, Jason A. Morrison, Mary Cathleen McKinney, Sean A. McKinney, Alejandro Sánchez Alvarado

## Abstract

Camera-type eyes are complex sensory organs susceptible to irreversible damage. Their repair is difficult to study due to the paucity of camera-type eye regeneration models. Identifying a genetically tractable organism with the ability to fully regenerate complete camera-type eyes would help overcome this difficulty. Here, we introduce the apple snail *Pomacea canaliculata*, capable of full regeneration of camera-type eyes even after complete resection. We defined anatomical components of *P. canaliculata* eyes and genes expressed during crucial steps of their regeneration. By exploiting the unique features of this organism, we successfully established the first stable mutant lines in apple snails. Our studies revealed that, akin to humans, *pax6* is indispensable for eye development in apple snails, establishing this as a research organism to unravel the mechanisms of camera-type eye regeneration. This work expands our understanding of complex sensory organ regeneration and offers new ways to explore this process.

## INTRODUCTION

The proper functioning of organs is intricately tied to their embryonic development, a complex and stereotypical multi-step process involving tissue organization and cell differentiation in the context of whole-body development. Post-embryogenesis, many animals lose the ability to fully regenerate new organs, some retain regenerative capacities within certain tissue or age and others exhibit remarkable abilities, regenerating entire organisms from small body portions^1,2^. Regeneration involves steps such as wound healing, detection of missing structures, activation of cell proliferation, determination of anatomical patterning, growth and integration of new tissues into the existing body^1,2^. Despite numerous studies on mechanisms driving regeneration in different organisms and tissues, many questions persist about the key features of a regeneration-permissive environment.

Eyes serve as fundamental sensory organs for numerous species, enabling exploration and interaction with their environments. The camera-type eye stands out as a complex and highly specialized organ, seemingly eluding complete regeneration after full resection resulting in irreversible loss of this organ after injury or damage. Camera-type eyes are capable of high-resolution image formation and are characterized by a single closed chamber, a lens, a cornea and a retina housing numerous photoreceptor cells equipped with molecular machinery for light detection. Examples of these can be found in human and more broadly in vertebrates, and among spiders and mollusks^3–5^. Camera-type eyes are just one of the eye types present among metazoans. The pigmented cup is a non-image-forming eye type, featured by planarians and distinguished by a cup housing photoreceptors and filled with dark pigments^3–5^. Bivalves, insects and decapods are examples of organisms with compound eyes, which result from several smaller units known as ocelli organized into a larger organ^3–5^.

In the past, research on eye regeneration has predominantly concentrated on complete regrowth of simpler planarian pigmented cups^6^ and on partial regeneration of camera-type eyes in vertebrates with robust regenerative abilities, such as fishes, newts and frogs. These remarkable animals can repair specific components following minor injuries, such as lens or retinal cell ablations, through the activation of the ciliary marginal zone or of Müller glial cells residing in the mature retina^7,8^. While these studies have significantly enhanced our understanding of eye regeneration, a mechanistic comprehension of complete adult camera-type eye regeneration remains elusive.

To overcome the existing limitation and expand our understanding of visual system regeneration, an organism characterized by camera-type eyes, robust regenerative potential and amenability to genome manipulation would be highly advantageous. The prevalence of camera-type eyes spans various animal phyla, including cnidarians, annelids, mollusks, crustaceans, arthropods and vertebrates^3–5^. In 1766, Spallanzani described the remarkable regenerative potential of garden snails following head amputations^9^. More recently, reports indicated the ability of certain gastropods to regenerate their visual systems^10–12^. These initial findings suggest gastropods could be useful organisms for investigating complete camera-type eye regeneration. Thus, we focused on developing a gastropod model amenable to genome manipulation^13^. *Pomacea canaliculata*, also known as the golden apple snail, is an amphibious freshwater gastropod native to South America and belonging to the *Ampullariidae* taxon^14^. These organisms exhibit resilience to diverse environmental conditions, breed throughout the year and have successfully completed their life cycle in captivity, making them an ideal organism for laboratory maintenance^14^. *P. canaliculata* is diploid^15^ and both its nuclear (440 Mb) and mitochondrial genomes have been sequenced, assembled and annotated^16–18^.

Here, we characterize the camera-type eye of *P. canaliculata* and describe the complete regeneration of this sensory organ after complete amputation. We also report on newly developed methods for collecting and micro-injecting zygotes and for culturing embryos. These protocols allowed us to introduce for the first time CRISPR-Cas9 technology to edit the apple snail genome and obtain stable mutant lines in *P. canaliculata.* Finally, we show that *pax6* function in eye development has been conserved in apple snails by developing the first *pax6*^-/-^ lophotrochozoan. This work opens the door to studying the function of genes potentially involved in camera-type eye regeneration in this, now genetically tractable, organism.

## RESULTS

### *P. canaliculata* has complex camera-type eyes

Camera-type eyes are present in vertebrates and characterized by one closed chamber with the light travelling through a transparent cornea, an anterior chamber, a lens, a posterior chamber and multiple retina layers to finally reach the apical part of the photoreceptors, also called the outer segment^3–5^. The outer segment of the photoreceptors is an area where the visual pigments, molecules that activate a signaling cascade when hit by photons, are hosted on the membranes^19,20^. Camera-type eyes can also be found in invertebrate organisms such as mollusks and specifically cephalopods as well as some gastropods, such as the tiger conch, *Conomurex luhuanus*^3–5,21^.

With the goal of finding an organism with camera-type eyes and high regenerative potential, we sought to determine if the apple snail *P. canaliculata* had camera-type eyes suitable for study (**Figure 1A**). The *P. canaliculata* eye bulb is located on top of a short eye stalk and the cornea is a circular transparent epithelium on its apical part (**Figure 1B**). Through micro-dissection, the retina with the enclosed lens can be isolated. The black pigmentation of the retina is uniformly distributed except for a circular area on its anterior side, which is not pigmented to allow for the transmission of the light (**Figure 1C**). The white optic nerve moves from the posterior side of the retina, through the eye stalk and reaches to the cerebral ganglia. The lens of *P. canaliculata* is oval and completely transparent and, through it, we can observe in-focus, sharp, inverted images, similar to the vertebrate lenses (**Figure 1D**).

**Figure 1.**
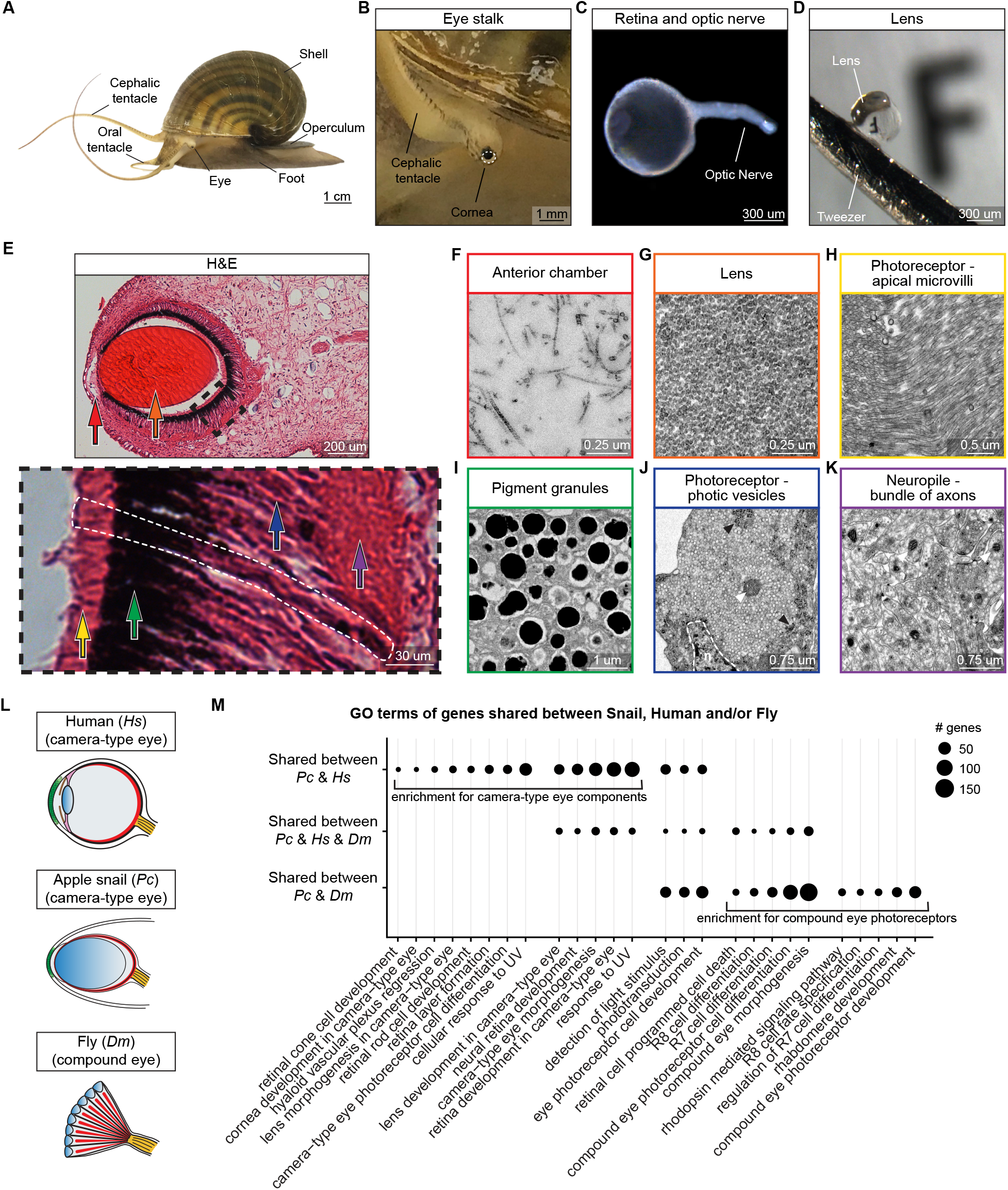
*P. canaliculata* has complex camera-type eyes. **(A)** Adult *P. canaliculata*. **(B)** Eye bulb and eye stalk with cornea (dashed line). **(C)** Isolated optic nerve and retina with the lens enclosed in it. **(D)** Isolated lens held with tweezers in front of a paper with the letter F printed in font size 5. **(E)** Hematoxylin and eosin (H&E) staining of *P. canaliculata* eye longitudinal sections. Cornea (dashed line on the left), anterior chamber (red arrow), lens (orange arrow), posterior chamber, retina, optic nerve and surrounding connective tissue, muscle tissue and extracellular matrix (ECM) can be distinguished. The inset highlights a photoreceptor (dashed line) and the retina layers: photoreceptor outer segments (yellow arrow), pigment granules (green arrow), photoreceptor inner segments (blue arrow) and neuropile (purple arrow). Arrows in (E) and images in (F-K) have the same color-code. TEM images of **(F)** anterior chamber filled with ECM; **(G)** lens with a densely packed structure; **(H)** outer segment of rhabdomeric photoreceptors characterized by microvilli; **(I)** pigment granules; **(J)** densely packed photic vesicles occupy an extensive portion of the photoreceptor cytoplasm together with rare mitochondria (white arrowhead) and ribosomes (black arrowheads), n=nucleus; **(K)** neuropile, or bundle of axons forming the optic nerve, filled with heterogeneous vesicles. **(L)** Schematic representation of eye anatomy for human (*Hs*), apple snail (*Pc*) and *Drosophila* (*Dm*). **(M)** Gene Ontology (GO) enrichment analysis of *P. canaliculata* (*Pc*) genes bioinformatically defined as orthologs of human (*Hs*) or fly (*Dm*) genes annotated with GO terms related to eye development and function. Adjusted *p* value cutoff of 1e-5. Manually selected representative terms (See Figure S1 and Table S1).

Longitudinal sections of the *P. canaliculata* eye show the presence of a cornea, an anterior chamber filled with loose extracellular matrix, a thin layer of unpigmented cells and the posterior chamber, delimited by the retina and mostly occupied by the lens (**Figure 1E, 1F and S1A**). Ultra-structurally, the lens is homogeneous and densely packed, but no structures resembling a cellular origin have been observed (**Figure 1G**), suggesting that *P. canaliculata* might develop its lens through a different strategy than vertebrates and cephalopods^22–24^.

The most abundant cells in the apple snail retina are rhabdomeric photoreceptors, a type of photoreceptor more prevalent among invertebrates and with microvilli in their outer segment (**Figure 1E, 1H and S1A-C**). The photoreceptor apices, or outer segments, face the lens, similar to cephalopods. A layer of cytoplasmic dark pigment granules is localized between the outer and inner segment of the photoreceptors, providing directional information and protection of deeper tissues from UV light (**Figure 1E and 1I**). The photoreceptor cytoplasm is filled with small and homogeneous photic vesicles and more rarely with mitochondria and ribosomes (**Figure 1J)**. The photoreceptor nuclei are basal, close to the neuropile or bundles of axons that move posteriorly to form the optic nerve (**Figure 1E**). In this basal region, we observed synapses between the basal part of the photoreceptors and axons, containing morphologically heterogeneous vesicles, larger and more electron-dense than the photic vesicles (**Figure 1K and S1H**). Additionally, we found several ciliated cells in multiple areas of the retina, such as intercalated in the microvilli and closer to the basal part of the photoreceptors (**Figure S1D-F**). Finally, the eye bulb is surrounded by connective tissue, muscle tissue and extracellular matrix (**Figure 1E and S1G**).

We conclude that *P. canaliculata* has complex camera-type eyes, that share some anatomical features with vertebrate eyes and others with camera-type eyes previously described in invertebrates, such as cephalopods and other gastropods.

### *P. canaliculata* eyes express conserved camera-type eye genes

Extensive comparative analyses performed among species support the hypothesis that similar anatomical structures of the visual system are the result of convergent evolution^25^. Concurrently, the pool of genes used for a camera-type eye development was determined to be conserved^26,27^. Genes well known to be expressed in vertebrate and cephalopod eye development include *pax6*, *vsx* and *rx*^28^. To analyze the genes expressed by apple snail eyes, we performed bulk RNA-Sequencing (RNA-Seq) on several *P. canaliculata* adult tissues, including the eye bulb and the isolated retina.

We first looked at the overlap between the genes present in *P. canaliculata* genome and those assigned to GO terms related to “visual system” in human (a vertebrate with camera-type eyes) and/or the fruit fly *Drosophila melanogaster* (an invertebrate with compound eyes) (**Figure 1L**). Interestingly, the group of genes shared between apple snails and humans is enriched with those involved in the overall morphogenesis of the camera-type eye and of its individual components, such as the development or formation of cornea, lens, photoreceptors and retina layers. The group of genes shared between apple snails and flies is enriched with those involved more specifically in the fly photoreceptor development. We observed genes shared between all the three animals to a lesser extent (**Figure 1M and Table S1**).

Additionally, we took advantage of the previously published vertebrate Gene Regulatory Networks (GRNs) driving optic cup patterning, camera-type eye morphogenesis and retina cell differentiation and looked for their *in silico* orthologs in *P. canaliculata* (**Figure S1I and S1J and Table S2**). We found 62% of these genes in the apple snail genome and for an additional 29% of these genes we found genes belonging to the broader gene family (**Table S2**).

Together, our analyses demonstrate that the apple snail genome and more specifically eye transcriptome includes several genes involved in the development of the vertebrate camera-type eye and its individual components. Additionally, *P. canaliculata* also has and expresses genes usually associated with *D. melanogaster* rhabdomeric photoreceptors. We can conclude that *P. canaliculata* shares molecular similarities with both vertebrate camera-type eyes and with invertebrate photoreceptors.

### *P. canaliculata* camera-type eyes can fully regenerate after complete amputation

To test if *P. canaliculata* can fully regenerate its camera-type eyes, we completely amputated the eye stalk, removing all eye bulb components (**Figure S2A**). Strikingly, we observed complete regeneration of the eye bulb and eye stalk within a month (**Figure 2A**). To characterize morphological changes and cell proliferation, we performed a time course of Hematoxylin and Eosin (H&E) staining as well as an anti-Phospho-Histone H3 (anti-H3P) antibody staining and 24 h BrdU pulse chase experiment.

**Figure 2.**
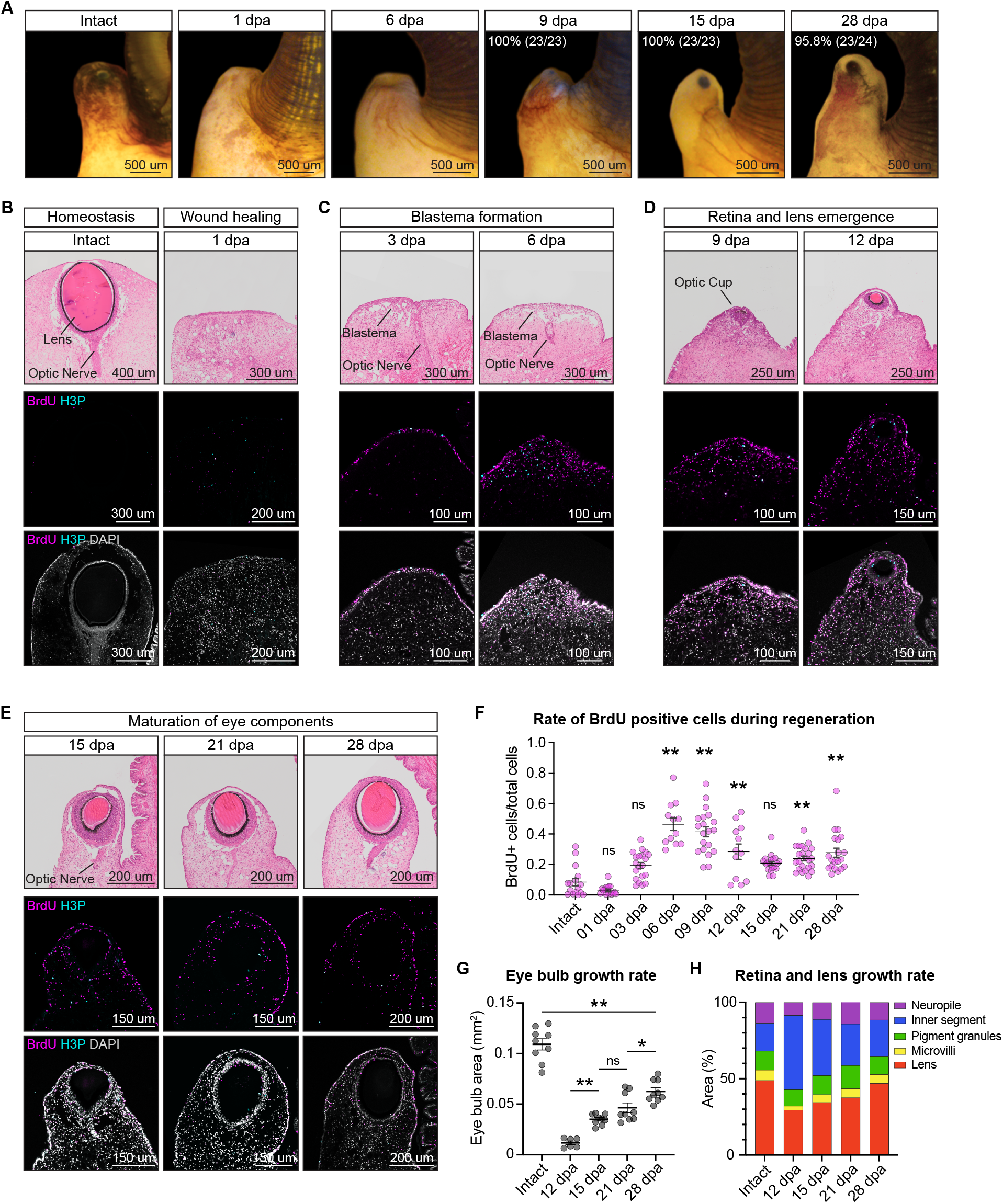
*P. canaliculata* camera-type eyes can fully regenerate after complete amputation. **(A)** Regeneration time course after complete amputation of the eye bulb. **(B-E)** Longitudinal sections of the eye stalk during eye regeneration time course stained with either H&E (top images) or BrdU and anti-H3P (central and bottom row of images). **(F)** Quantification of BrdU positive cells in the regenerating tissue area during the eye regeneration time course. n = 3 to 6 snails/time point, 4 sections each. **(G)** Quantification of the eye bulb area during the eye regeneration time course. n = 3 snails/time point, 3 sections each. **(H)** Quantification of the relative area of the eye bulb components (lens, photoreceptor microvilli, pigmented layer, photoreceptor inner segment and neuropile) during the eye regeneration time course. Data are represented as mean ± SEM. n = 3 snails/time point, 3 sections each. * = *p* value < 0.05, ** = *p* value < 0.01, ns = non-significant (see Figure S2).

We identified four main stages of eye regeneration in *P. canaliculata*: wound healing, blastema formation, retina and lens emergence and eye components maturation. The wound completely heals in the first 24 h, concurrent with a significant increase of H3P positive cells (**Figure 2B, S2B and S2C**). The blastema, tissue composed of loose proliferating cells and developed by several animals during the regeneration process^1,2^, forms at 3 and 6 days post amputation (dpa) (**Figure 2C, 2F, S2B and S2C**). The new optic cup, invagination of retinal cell precursors organized in a cup that closes on it-self, develops at 9 dpa. This optic cup is localized close to the apical portion of the eye stalk and already presents some black pigmentation (**Figure 2D**). At 12 dpa the regenerating retina is well defined with the lens developing inside it (**Figure 2D**). All the major morphological features of the eye bulb are present by 15 dpa, including the optic nerve (**Figure 2E**). At later time points (21 and 28 dpa) all these anatomical components continue to mature and grow (**Figure 2E**).

We determined the growth rate of the regenerating eye by calculating the eye bulb area and found that at 28 dpa it is still significantly smaller than the one of the original eye (**Figure 2G**). Interestingly, we observed the different components (lens, photoreceptor microvilli, pigment granules, photoreceptor inner segment and neuropile) do not form simultaneously and do not grow at the same rate. Specifically, the photoreceptor inner segment occupies a significant percentage of the eye bulb at 12 and 15 dpa and grows less over time compared to the other layers. By 28 dpa, the relative sizes of these components become similar to those observed in the intact eyes (**Figure 2H and S2D-H**).

Altogether, these data show that *P. canaliculata* fully regenerates its camera-type eyes, healing the wound in the first 24 h and forming a blastema from 3 to 6 dpa. Intense cell proliferation accompanies retina emergence through the formation of an optic cup. After lens emergence at 12 dpa, there is a phase of maturation and growth of the eye components through cell proliferation and retinal layer differentiation.

### *P. canaliculata* regenerating eyes express genes driving vertebrate eye development

Following our morphological characterization, we next wanted to evaluate the changes in gene expression during *P. canaliculata* eye regeneration. We generated bulk transcriptomes at morphologically defined regeneration time points (**Figure 3A and S3A**). As expected, the segregation of the samples in the MDS plot mirrors morphology showing that the early time points of regeneration (1 and 3 dpa) are transcriptionally the most different from the intact eyes (largest distance in Dimension 1) while the later time points (21 and 28 dpa) are the closest to the intact eyes (**Figure 3B**). All time points are chronologically positioned in an arc shape from 1 to 28 dpa with the peak at 6 dpa (highest distance in Dimension 2) (**Figure 3B**). This distribution of the samples leads us to hypothesize that the genes homeostatically expressed in the eye components that are lost at 1 dpa and slowly regenerated at the various time points, are likely a major driver in Dimension 1. Similarly, Dimension 2 might mainly reflect genes related to cell division and cell proliferation that are enriched in the blastema (3, 6 and 9 dpa).

**Figure 3.**
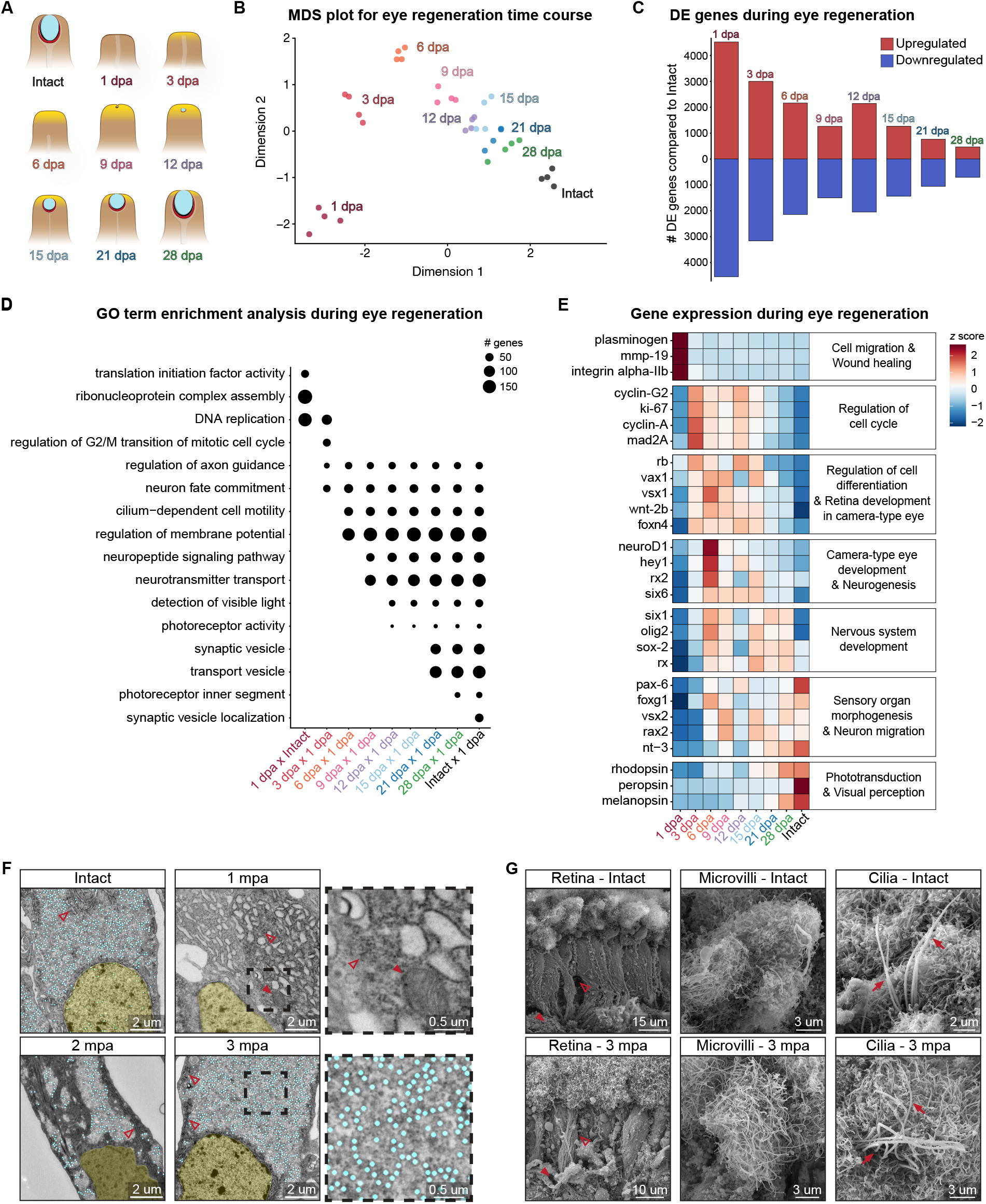
*P. canaliculata* regenerating eyes express genes driving vertebrate eye development. **(A)** Schematic representation of the main morphological defined time points during eye regeneration. Blastema/cell proliferation (yellow), lens (cyan), retina (red). **(B)** MDS plot of RNA-Seq samples collected during eye regeneration time course. Dimension 1 explains 41.8% of variance and Dimension 2 13.72%. n = 4 samples/time point (see Figure S3). **(C)** Number of differentially expressed (DE) genes (up-regulated in red and down-regulated in blue) compared to Intact samples [Log Fold Change (LogFC) >= 0 and FDR <= 1e-5]. **(D)** Gene Ontology (GO) enrichment analysis using significantly up-regulated genes (LogFC >= 0 and FDR <= 1e-5) in 1 day post injection (dpa) vs Intact and in 3, 6, 9, 12, 15, 21 and 28 dpa vs 1 dpa comparisons to highlight the features recovered throughout regeneration. GO terms used in this representation were manually selected from GO enrichment analysis run on the up-regulated genes obtained from comparing each time point with the previous one (*p* value <= 0.01, q value <= 0.05) (see Figure S4 and Table S3). **(E)** Heatmap of gene expression changes during eye regeneration time course. These genes represent the main stages of eye regeneration, and many of them are known, from previous studies, to be involved in vertebrate eye development (*z* scores calculated from TPMs). Gene categories on the right of the heatmap are based on GO annotations. **(F)** TEM images of the photoreceptor cytoplasm in the retina of Intact, 1, 2 and 3 months post amputation (mpa) eyes. Nuclei (yellow), photic vesicles (cyan), ribosomes and rough endoplasmic reticulum (empty arrowhead) and mitochondria (full arrowhead). **(G)** SEM images of the retina, photoreceptor microvilli and apical cilia observed in Intact or 3 mpa eyes. At both time points, the retina is constituted by long photoreceptors (empty arrowhead), neuropile (on the bottom, full arrowhead) and microvilli (on the top). The photoreceptor microvilli at 3 mpa show an apical flat and wide expansion of the membrane like the Intact eyes. Cilia (full arrow) have been observed intercalated in the photoreceptor microvilli both in Intact and 3 mpa retina.

To further dissect the molecular dynamics of snail eye regeneration, we calculated the number of differentially expressed (DE) genes between the regeneration time points and the intact eye. While initially there are about 9000 DE genes, this number steadily decreases until 28 dpa where there are only 468 genes up-regulated and 707 genes down-regulated compared to the intact eye (**Figure 3C, S3B and S3C**). To gain a better understanding of the steps followed by the regeneration process, GO term enrichment analysis and heatmaps were performed on DE genes (**Figure 3D, 3E and S4A-F and Table S3**). We found an enrichment of genes involved in wound healing and cell migration in the first 24 h after amputation. Genes related to DNA replication and regulation of the cell cycle were up-regulated at 3 dpa and persisted throughout the regeneration time points. At 3 and 6 dpa genes associated with neurogenesis, axon guidance and camera-type eye development are already enriched and up-regulated. Enrichment for the detection of visible light and photoreceptor activities starts at 12 dpa, while a molecular signature for phototransduction and visual perception happens later in the time course. Interestingly, terms that are enriched and expressed at a higher level in the intact eye compared to the 28 dpa are related to synaptic vesicle biology and visual pigments (**Figure 3D, 3E and S4A-F and Table S3**). This difference suggests the regenerating eye may require a longer time to fully complete the maturation of certain cell types.

To identify ultra-structural differences among the intact and the regenerating eyes, we analyzed the regenerating tissue at longer time points with electron microscopy. Interestingly, the main differences that we noticed throughout the time course were related to the presence of photic vesicles. At 1 month post amputation (mpa) the cytoplasm of the photoreceptors does not contain any photic vesicles, while there is abundance of ribosomes and endoplasmic reticulum (ER). Some areas with photic vesicles were found in the 2 mpa tissue, although less packed and surrounded by areas still containing ribosomes and ER. By 3 mpa the photic vesicle distribution is indistinguishable from the intact eye (**Figure 3F**). At this time point we also observed that the microvilli of the rhabdomeric photoreceptors and the cilia among the microvilli were recovered (**Figure 3G**).

Overall, these data provide a molecular blueprint of snail eye regeneration, as well as the molecular and ultra-structural differences still present between the 28 dpa and the intact eye.

### *P. canaliculata* zygotes can be cultured *ex ovo* and micro-injected with exogenous mRNA

To test gene function and investigate the molecular mechanisms driving development and regeneration of eyes in *P. canaliculata*, we sought to build tools to genetically modify these snails. Since the oocytes are internally fertilized, we developed a protocol to efficiently collect zygotes, opening the external capsules as soon as they are laid by the female. These embryos cannot fully develop without the pink perivitelline fluid (PVF) present inside the capsule and representing their main source of nutrients until hatching. To grow embryos after manipulations, we developed a protocol to culture them *ex ovo* in an extract of perivitelline fluid (ePVF) (**Figure 4A**). To avoid evaporation, drops of ePVF were placed in a Petri dish and covered in paraffin oil. The embryos cultured *ex ovo* complete development in 10-13 days, similar to embryos growing in their original capsules (**Figure 4B and S5A**). In addition to growing manipulated embryos that were removed from their capsules, this protocol now allows us, for the first time, to observe embryonic development live (**Figure 4B and S5A**).

**Figure 4.**
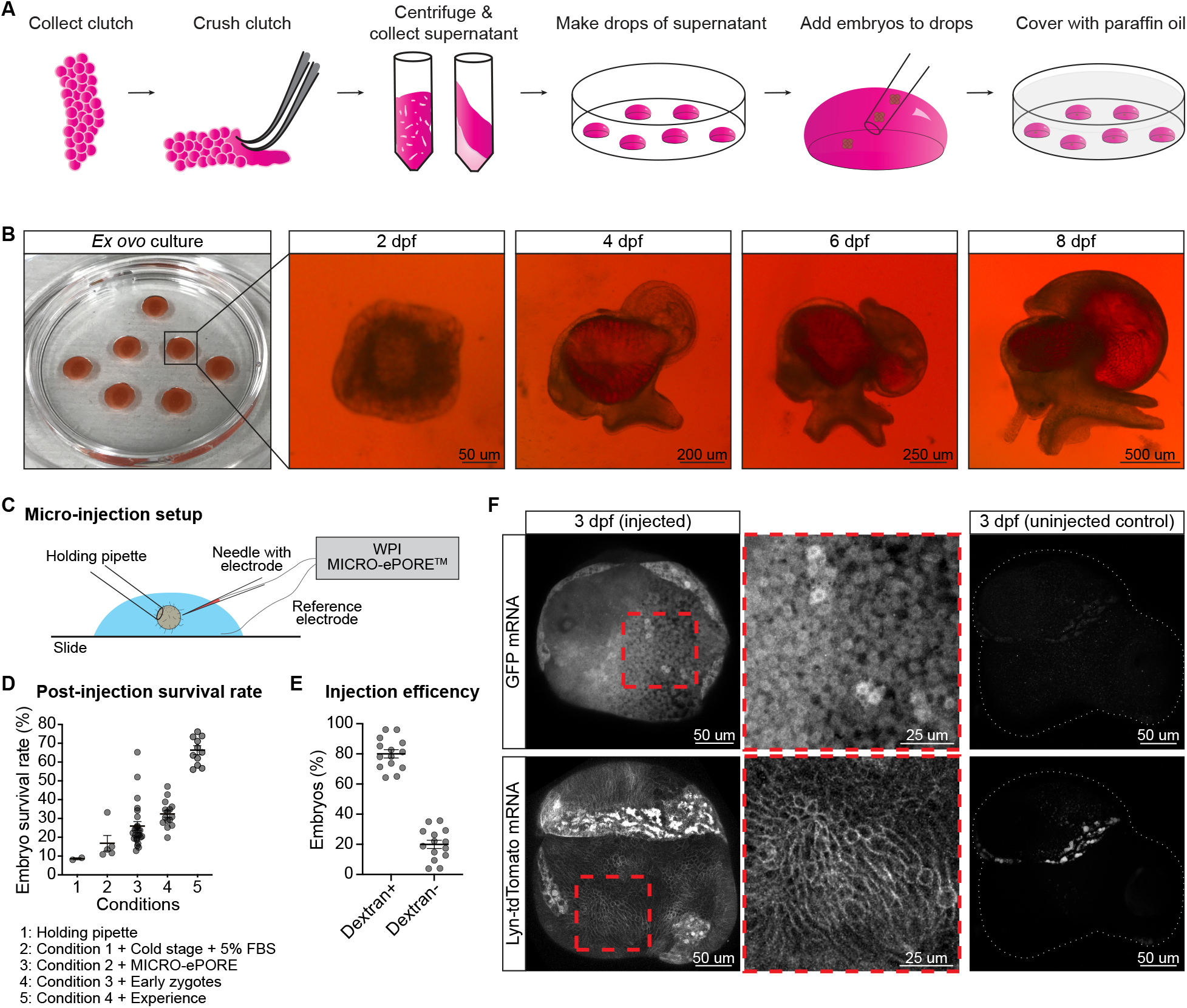
*P. canaliculata* zygotes can be collected, micro-injected with exogenous mRNA and cultured *ex ovo*. **(A)** Schematic representation of *ex ovo* culture protocol steps. Clutches are collected, crushed and centrifuged. The supernatant is collected and used to make drops where the embryos can be cultured. Everything is covered with paraffin oil to avoid evaporation. **(B)** Images of embryos cultured at 0 days post fertilization (dpf) and developing *ex ovo*. **(C)** Schematic representation of the setup used for *P. canaliculata* zygote micro-injection (see Figure S5). **(D)** Embryonic survival rate after micro-injection using different micro-injection protocols. Each dot represents an injection session. **(E)** Percentage of embryos Dextran-positive, used as read-out of successfully injected embryos. Each dot represents an injection session. **(F)** Confocal images of 3 dpf embryos injected with GFP mRNA or Lyn-tdTomato mRNA. GFP localizes in the cytoplasm, the membrane targeted tdTomato localizes in the cell membranes and the uninjected controls show some autofluorescence with a stereotypical localization in the red channel. Data are represented as mean ± SEM.

As apple snail zygotes are similar in size to one-cell mouse embryos (approximately 100 um in diameter), we adopted a modified micro-injection system similar to the one used for creation of transgenic mice. This system included the standard inverted microscope, a glass micro-injection needle and a glass holding pipette. In addition to this equipment, we incorporated the use of a cold microscope stage and a MICRO-ePORE pinpoint electroporation system (**Figure 4C, S5B and S5C**). Multiple rounds of micro-injection were performed to further optimize equipment and reagents resulting in increasing survival rates. Using a holding pipette, a cold stage (at 12 °C), media supplemented with FBS and the MICRO-ePORE, we reached an average of 30-40% survival rate that increased to more than 65% with user experience (**Figure 4D**). Initially, we micro-injected fluorescent dextran and we determined that an average of 80% of the embryos which survived showed fluorescent signal, confirming a successful micro-injection (**Figure 4E**).

Using the protocols we developed to micro-inject and culture *P. canaliculata* embryos, we next asked if they could translate exogenous mRNA. After injecting the embryos with mRNA for either GFP or membrane targeted tdTomato, we observed fluorescence in the correct wavelength and subcellular localization (**Figure 4F**). In both cases the fluorescent signal was detectable up to 4 days post injection (dpi).

Overall, we developed tools and approaches that are crucial for making apple snails amenable for a broad variety of experimental designs. Embryos can be collected at any stage of interest and then cultured *ex ovo*, where it is possible to observe and image them over time. Moreover, we optimized micro-injections in zygotes and demonstrated that exogenous mRNA can be used as strategy for gene over-expression or lineage tracing experiments. Together these represent fundamental steps toward bringing robust approaches for genetic manipulations in apple snails.

### *P. canaliculata* has a *pax6* gene highly expressed in embryonic eye buds

*pax6* is a regulatory gene with a fundamental role in brain and eye development. In mouse and *D. melanogaster*, *pax6* expression is required for proper eye formation^29,30^. The *P. canaliculata* genome^16,17^ has 5 genes annotated as *pax6-like* (NCBI RefSeq GCF_003073045.1). To determine if these genes contain both domains characteristic of *pax6* (PAX and HOX domains) and are similar to the human and fly proteins, we annotated the domains, predicted the 3D folding and evaluated the amino acid conservation for all the sequences.

We found that two out of the five genes identified in apple snails have both PAX and HOX domains, but only one (LOC112559942) has a complete PAX domain and a much higher amino acid conservation than the others (**Figure 5A, S6A and S6B and Table S4**). This result was confirmed by running a phylogenetic relationship and analyzing their expression levels. The gene LOC112559942 is phylogenetically closer to human fly *pax6* and is more highly expressed in the adult retina than the other candidate genes (**Figure S6C and S6D**). This suggests that LOC112559942 is an ortholog of the human and fly *pax6*.

**Figure 5.**
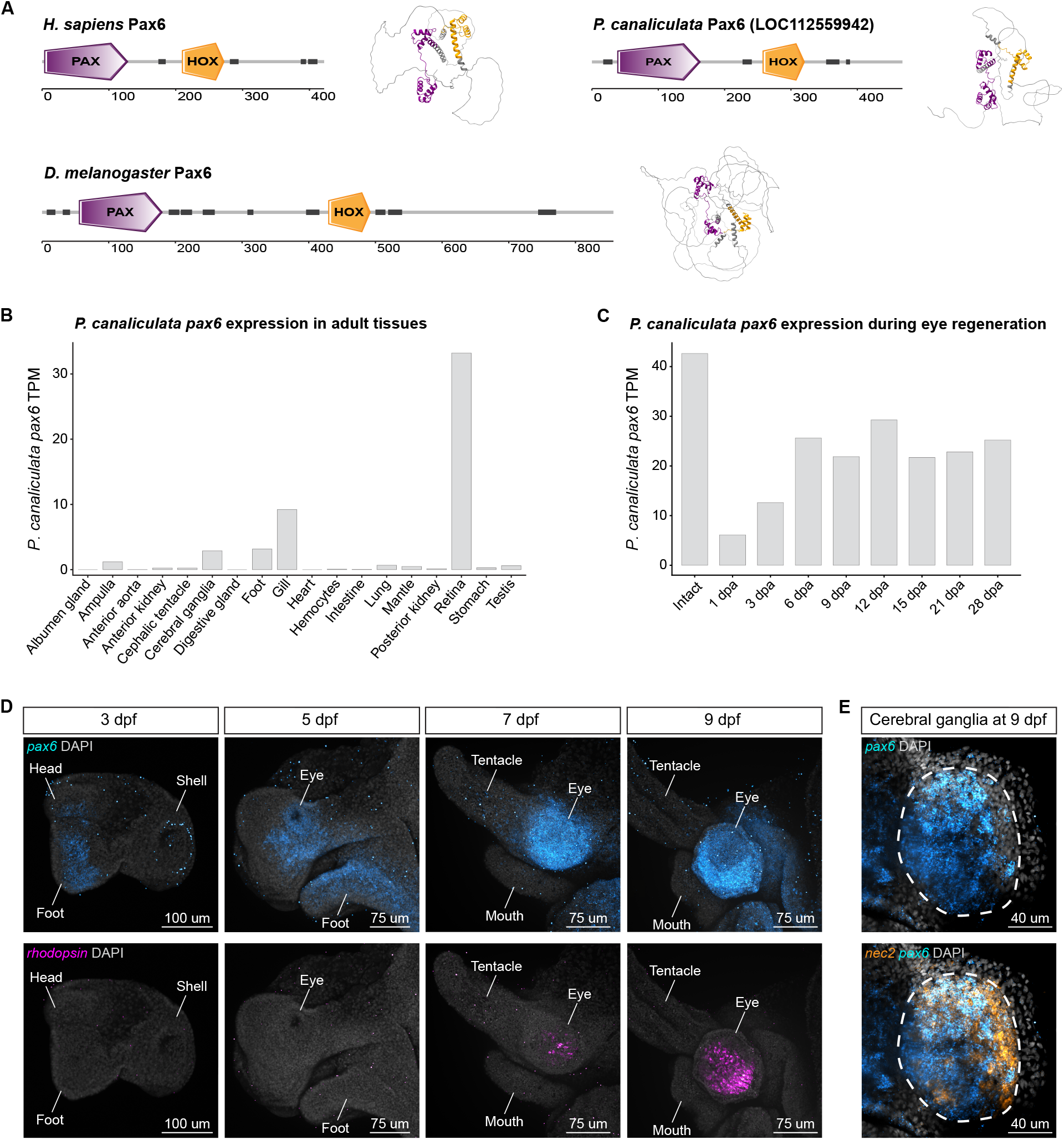
*P. canaliculata* has a *pax6* gene highly expressed in the eye buds. **(A)** Domain and 3D-folding predictions for human, *D. melanogaster* and *P. canaliculata pax6* genes. Purple = PAX domain, orange = HOX domain, dark gray = disordered regions (see Figure S6). *pax6* expression levels **(B)** in isolated adult tissues and **(C)** during eye regeneration, expressed as TPMs. **(D)** Confocal images of *pax6* and *rhodopsin* HCR during embryonic development. *rhodopsin* highlights the localization of the retina and the differentiation of the photoreceptors. **(E)** Confocal images of *pax6* and *nec2* HCR during embryonic development. *nec2*, also known as *proprotein convertase 2* (*pc2*) or *prohormone convertase 2*, has been used as marker for the nervous system.

To gain a better understanding of *pax6* expression and dynamics, we evaluated its expression through RNA-Seq analysis across multiple adult tissue, during eye regeneration time course and HCR *in situ* hybridization during embryonic development. Transcriptomic analysis shows that the retina is the adult tissue where *pax6* is more highly expressed and during complete eye regeneration pax6 expression increases from 1 to 6 dpa (**Figure 5B and 5C**). Since *P. canaliculata* is characterized by direct development (*i.e.*, lack of metamorphosis or free-living larval stage) we performed HCR on developing embryos and mapped *pax6* expression to the embryonic equivalent of adult organs. HCR during embryo development shows that at 3 days post fertilization (dpf) *pax6* is expressed in the anterior part of the embryos including the cephalic area and the foot. At 5 dpf fluorescent signal can be observed in the eye buds, anteriorly to the eye buds and in the foot. The *pax6* expression seems more clearly confined in the eye stalk at 7 dpf and 9 dpf. At these stages, we can also see the expression of rhodopsin, a marker for photoreceptor cells, that are differentiating in the embryonic retina (**Figure 5D**). In 9 dpf embryos we also observed the *pax6* expression in the cerebral ganglia together with the neuronal marker, *nec2* (**Figure 5E**).

In summary, we found a *pax6* gene in *P. canaliculata* that structurally resembles the human and *D. melanogaster pax6* genes and is expressed in the apple snail eyes, both during development and in adults.

### CRISPR mutant snails reveal *P. canaliculata pax6* is required for eye formation

To test pax6 function in *P. canaliculata* and evaluate if its role in eye development has been conserved, we applied the CRISPR-Cas9 system in apple snails. Taking advantage of the new protocols we developed for micro-injection and *ex ovo* culture, we established the protocol to generate loss of function mutations through micro-injection of guideRNA (gRNA) and Cas9 protein (**Figure S7A, S7B and S7C**). To ensure reproducibility and robustness of our protocol for gene knock-out in apple snails, we targeted three different genes (*pax6*, *dia2* and *pitx*) and we successfully generated mutant lines for all of them (**Figure 6A-D and S7D-I and Table S5**).

**Figure 6.**
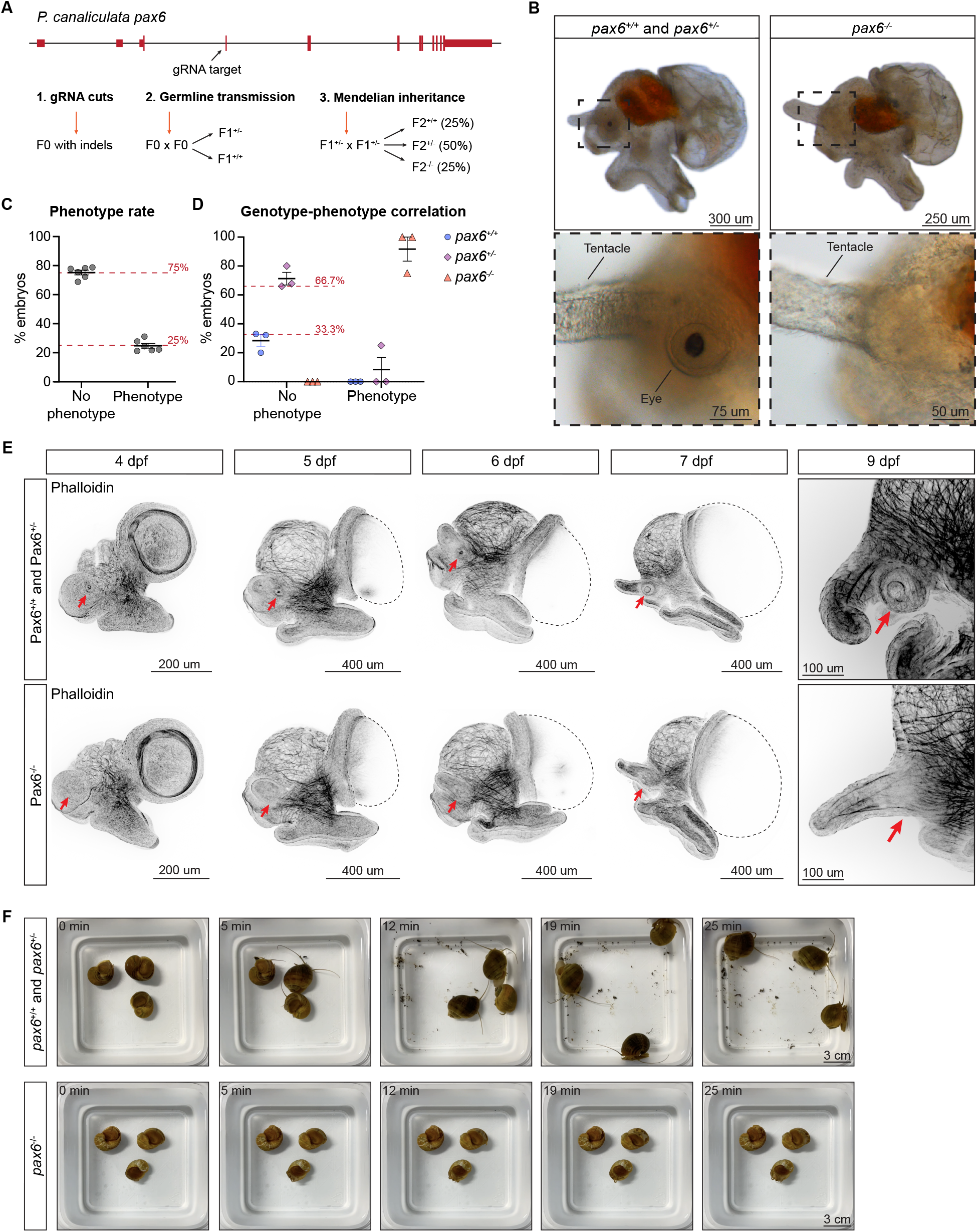
CRISPR mutant snails reveal *P. canaliculata pax6* is required for eye formation. **(A)** Schematic representation of the *P. canaliculata pax6* genomic region, the gRNA target site, and the crossing strategy used to establish a stable mutant line and to obtain homozygous mutant animals (see Figure S7 and Table S5). **(B)** Images of wild-type and pax6^-/-^ 9 dpf embryos and insets of the region at the base of the cephalic tentacle. Wild-type and pax6^+/-^ animals have eyes while the pax6^-/-^ snails lack all the eye bulb and eye stalk components. **(C)** The percentage of F2 embryos with the “lack of eye” phenotype correspond to the expected 25%. n = 6 clutches. **(D)** Correlation between genotypes and phenotypes in F2 embryos shows that the lack of eyes corresponds to homozygous mutants. n = 3 clutches. **(E)** Confocal images of wild-type and pax6^-/-^ embryos stained with phalloidin during a developmental time course. Phalloidin shows that at 4 dpf there is a first morphological evidence of eye formation through cell wall contraction in the cephalic area of the embryos. The red arrows point at the area where the eye develops (in wild-type and pax6^+/-^ snails) or where it should be developing (in pax6^-/-^ snails). **(F)** Still images of time lapses showing adult wild-type and pax6^-/-^ snails free to move in their tanks (see Video S1). Data are represented as mean ± SEM.

Specifically for *pax6,* a gRNA was designed to target Exon 3 out of the 11 exons present in the gene. F0 individuals were grown to adults and crossed with each other or wild-type animals (**Figure 6A and S7D-F**). The F1s were then genotyped to test for germline transmission of the mutation in the targeted site (**Figure 6A and S7G**) and the animals heterozygous for the mutation were incrossed to generate F2s or outcrossed to maintain the line. Genotypes of F2 siblings confirmed that the mutation was distributed among the offspring following Mendelian inheritance (**Figure 6A and S7H and S7I**).

The F2 embryos obtained from the *pax6* mutant line were then used to analyze the phenotype. Homozygous mutant embryos at 9 dpf do not have retina nor eye stalk. Close to the tentacle, there is no resemblance of any eye structure, while all the other cephalic organs do not present any observable phenotype (**Figure 6B**). The phenotype was observed in about 25% of the F2 embryos and subsequent genotype of these animals showed that only homozygous mutant embryos presented the phenotype (**Figure 6C and 6D**). Staining with phalloidin during an embryonic development time course highlighted that in absence of *pax6* there is no sign of eye formation at any moment of embryogenesis (**Figure 6E**). The earliest time point where eye formation is detected in wild-type embryos is 4 dpf as an area of higher condensation of cell membrane in the cephalic region close to the tentacle buds. Mutant embryos at 4 dpf or later do not show the same cellular dynamics and the epithelial cells close to the tentacle buds are homogenously distributed (**Figure 6E**).

Additionally, a behavioral phenotype was observed in *pax6* mutants, from hatchings to adults. Wild-type and heterozygous snails, from the time they hatch, are significantly active and can easily flip on to their foot to move around and explore their environment. Both while on their dorsal side and while moving on their foot, they spread their cephalic tentacles and move them around. Homozygous mutant snails, on the other side, continuously lay on their dorsal side at the bottom of the tank, are unable to rotate on their ventral side to move using their foot and keep their cephalic tentacles constantly curled close to the head. Wild-type adult snails whose eyes were amputated did not phenocopy the homozygous mutant behavior and the way they move around and use their tentacles was indistinguishable from the wild-types (**Figure 6F and Video S1**). This suggests that the lack of eyes is not the cause of the observed behavioral phenotype, implying that *pax6* is necessary, not only for eye formation, but also for proper development of components that are crucial for locomotion, for the use of cephalic tentacles and foot and maybe, even for the gravity perception.

This is the first time that a *pax6* stable knock-out line has been developed in a lophotrochozoan confirming, through loss of function, the conserved role of this transcription factor in eye development. We showed that at least one copy of *pax6* is necessary to activate the eye development program and that *pax6* role during brain development might be conserved in apple snails. Overall, the data presented here show that *P. canaliculata* is now a genetically tractable research organism that can be adopted to study gene function in biological processes of interest.

## DISCUSSION

Our eyes are complex and delicate organs that can only rarely recover from damage or aging. Among research organisms, some vertebrates, like zebrafish, salamanders and frogs, can recover from minor injuries to specific retinal cell types or individual eye components^7,8^. Meanwhile, some invertebrates, like planarians, can fully regenerate their eyes, but their visual system has a very different and more simple anatomical structure^6^. Our work aimed to bridge this gap by identifying an organism capable of regenerating camera-type eyes, which although classically associated with vertebrates, can also be found in some invertebrates^3–5^. Here, we report the discovery that the camera-type eyes of the freshwater apple snail *P. canaliculata* can fully regenerate. We also developed protocols and tools to demonstrate the amenability of this novel regeneration organism to genetic manipulation. These resources not only open the door to answering fundamental questions about camera-type eye regeneration but also make *P. canaliculata* an innovative system where numerous other aspects of biology can be explored, since this is one of the very few mollusks where stable mutant lines can be established^31^.

### Animal visual systems: anatomical similarities and differences

The visual system is important for many aspects of animal physiology, such as navigating the environment, mating and regulating activities throughout the day. The eyes are one of the main components of the visual system and they can be classified based on their anatomical structure, components and light path^3–5^. While vertebrates have camera-type eyes, invertebrates show a broader spectrum of eye types^3–5^. The taxon *Mollusca* includes animals with extremely different body plans and lifestyles and almost all the known eye types are represented in this group^3–5^. This diversity offers the potential to answer questions about evolution of the visual system as well as the relationship between the different eye types.

Here, we show that the gastropod *P. canaliculata* has camera-type eyes. Interestingly, the animal environment and behavior are correlated to specific properties of some eye components, such as cornea and lens^3–5^. The human lens is biconvex, aquatic animals, such as zebrafish, have a spherical lens, and nocturnal animals, like mice, usually have larger lenses^3–5^. Apple snails are aquatic and relatively more active during the night and their lenses are round and large, key features usually associated to these traits.

Vertebrates have an inverted retina, with the apical part or outer segment of the photoreceptor pointing towards the external part of the eye^5^. These photoreceptors (rods and cones in humans) are defined as ciliary because of a modified cilium present between the inner segment, more basal with the nuclei and other organelles, and the outer segment, constituted by membrane discs and folds to host the visual pigments, such as opsins^19,32^. Like cephalopods and other invertebrates with camera-type eyes^5,33^, *P. canaliculata* retina is not inverted and thus the outer segment of the photoreceptor points towards the lens. The pigmented layer in the retina contains dark pigments that creates a physical barrier to protect deeper tissue from the light and provide directional information about the light source^3–5^. Interestingly, the pigmented layers of both vertebrates and *P. canaliculata*, although differently positioned relative to the retina, are localized immediately behind the outer segment of the photoreceptor, highlighting conserved function.

The photoreceptor anatomy, their visual pigments and their signal transduction pathways have a significant role in determining what the animals can see and consequently are strongly correlated with animal behaviors. Most photoreceptors found in invertebrates, independently from the eye type, are rhabdomeric photoreceptors. These photoreceptors do not have a modified cilium and their outer segment is constituted by microvilli instead of stacked membranes^19,32^. Here, we show that *P. canaliculata* has rhabdomeric photoreceptors. However, we also observed cells with centrioles and/or cilia and detected the expression of rod and cone-specific genes. Although additional data needs to be collected in apple snails, recent examples of invertebrates with both ciliary and rhabdomeric photoreceptors have been reported^19^.

*P. canaliculata* camera-type eyes bring together multiple anatomical and molecular features from both vertebrate and invertebrate visual systems. This hybrid anatomy was previously observed in cephalopods, highlighting how historical classifications and definitions might need to be revisited. Both apple snails and cephalopods are of great interest to better understand the evolutionary development of camera-type eyes and, more in general, of visual systems among invertebrates.

### Regeneration of missing body parts and specifically eyes

Regeneration is the process of replacing damaged or lost body parts. Regeneration can happen through different mechanisms; the more studied mechanisms are morphallaxis (reorganization of pre-existing tissue) and epimorphosis (intense cell proliferation and blastema formation)^1,2,34^. The regeneration of sensory organs, such as the eyes, represents a unique paradigm to study, considering their important function and their complex anatomy. Eyes are composed of multiple structures perfectly tuned with each other, with relevant neuronal components that are precisely wired to the brain for information integration and interpretation^35^. Though axolotls can regenerate the lens and zebrafish can regenerate retina cell types, there are no known examples of vertebrates that can fully regenerate their eyes after complete resection^7,8^. Lens and retina regeneration in axolotls and in zebrafish happen through morphallaxis (no blastema formation) with specific cell types in well-defined regions of the eye, such as Müller glia in the ciliary marginal zone, transdifferentiating, proliferating and replacing the lost cell types^7,8^.

Lophotrochozoans, and specifically gastropods, are known for their regeneration potential, with some more extreme cases of entire eye or even entire head regeneration^9–12,36,37^. Here, we show that adult and sexually mature *P. canaliculata* can regenerate its camera-type eyes after full amputation. This is a remarkable characteristic, since other organisms, like the snail *Achantina fulica*, lose or significantly decrease their regeneration abilities with age^12^. *P. canaliculata* eye regeneration happens through epimorphosis, with intense cell proliferation and formation of a blastema. The cellular and molecular mechanisms used for this regeneration remain unknown. In many other invertebrates with high regeneration potential through epimorphosis, a stem cell population that is activated after injury has been identified^2,34,38,39^. We speculate that apple snails also have a stem cell population responsible for the formation of the blastema, although additional experiments are required. However, the potential involvement of cell dedifferentiation or transdifferentiation in this process cannot be excluded.

Finally, previous evidence underlines the importance of both the immune system and the nervous system in creating regeneration permissive niches and supporting the new tissue growth. Müller glia cells have been shown to be necessary for retinal cells regeneration in zebrafish. In apple snails, it has been previously observed that clodronate treatment depletes circulating phagocytic hemocytes and delays the formation of the blastema after the amputation of the cephalic tentacle^40^. This suggests that the roles of the immune and nervous systems in *P. canaliculata* eye regeneration need to be further investigated.

Altogether, *P. canaliculata* is an extraordinary organism that brings together the presence of camera-type eyes and a high regenerative potential in adults. This represents a unique opportunity to study new aspects of complex sensory organ regeneration.

### Established and emerging organisms in research

Currently established research organisms have been selected in the past for practical reasons such as ease to maintain in captivity and year-round and frequent breeding that produces a considerable number of offspring. These animals also have a relatively fast generation time and some particularly interesting or unique features that made them essential to advance specific fields (*e.g.*, transparent zebrafish embryos or low chromosome number in *D. melanogaster*)^41,42^. Because of their genetics and widespread use, these were among the first animals in which methods to test gene function were developed and whose genomes were sequenced^43,44^. Recently, technological advances such as the reduced cost of sequencing and the advent of CRISPR systems have allowed the establishment of new research organisms^45–52^.

The freshwater apple snail *P. canaliculata* is a resilient animal that can complete its life cycle in an aquarium and breed year-round providing hundreds of embryos per clutch^14^. Apple snails are diploid^15^ and their genome is non-duplicated and smaller than many other mollusks^31,53^, which facilitated its sequencing^16,17^. These are important features for establishing *P. canaliculata* as a new genetically tractable organism. Here, we showed how we overcame challenges related to collecting and micro-injecting zygotes, raising F0s to adults and genotyping animals to establish the first stable mutant lines of apple snails using the CRISPR-Cas9 technology. Moreover, multiple mutant lines were established to confirm that this was a robust and reproducible protocol.

*P. canaliculata* has many features that make it a system with potential to be extensively used in the laboratory and it also has the remarkable and rare feature of being able to fully regenerate camera-type eyes, making it a system of great value to uncover the mechanisms of complex sensory system development and regeneration. This work provides fundamental tools which open avenues for the scientific inquiry of molluscan and lophotrochozoan biology.

### Shared regulatory network for eye development

In the past few decades, many studies were performed to gain a better understanding of the GRNs controlling eye morphogenesis and specific cell type differentiation^28,54^. The transcription factor *pax6* has been reported in multiple species, with and without camera-type eyes, as one of the initiators of eye development. Studies in *Drosophila* demonstrated that *pax6* expression is both necessary and sufficient for eye development^55^. In vertebrates, optic vesicles and optic cup determination happens through a signaling cascade activated by pax6 and involving several other eye field transcription factors^28,56^. Once the eye field has been determined and organized into specific regions, the neural retina progenitors start expressing genes downstream of *pax6* that specify the retinal cell types^28^. *P. canaliculata* has most of these genes in its genome and many are expressed in the adult eye bulb. This provides an opportunity to compare and contrast differences in the GRNs for eye development in between *P. canaliculata* and other models. For example, *P. canaliculata* pax6 has well conserved sequence, domains and 3D structure when compared to human PAX6 and it is expressed in both adults and embryonic eye. Pax6 is known to be involved not only in development of the eyes but also of the brain, nose, pancreas and pituitary gland^30,57^. Previous work on lophotrochozoans showed the presence of a *pax6* gene and its expression related to eye and brain development^6,58,59^. Here, we knocked-out *pax6* for the first time in a lophotrochozoan, showing *pax6* is necessary for eye development and likely plays a role in the development of cerebral ganglia. This suggests a conserved function for *pax6* between vertebrates and apple snails. We can extend this type of analysis to other components of the GRN.

So far, we explored the function of *pax6* gene during embryonic development. Based on our transcriptome analysis, where *pax6* is expressed in the early steps of regeneration, we hypothesized that this gene also plays a crucial role in adult eye regeneration. Future experiments will test the role of *pax6* in adults, as well as define the GRN to which *pax6* belongs providing exciting information about the mechanisms that control and allow complete regeneration of camera-type eyes in *P. canaliculata*. The experiments we have conducted illustrate how apple snails are an excellent model system poised for utilizing a broad range of experimental approaches to gain mechanistic insights into the evolution, development and regeneration of camera type eyes.

## ACKNOWLEDGEMENTS

We would like to thank Julia Peloggia de Castro and Drs. Robb Krumlauf, Stephanie Nowotarski, Frederick Mann and Viraj Doddihal for critical reading of the manuscript. We also thank the Sánchez Alvarado lab members for insightful discussions; Tonyea Inglis for the support in applying for the apple snail permit; the Stowers Institute for Medical Research (SIMR) Invertebrates Core, Carlos Barradas Chacón and Robert Schnittker for maintaining the *P. canaliculata* colony; Davide Malagoli for providing animals to establish the SIMR apple snail colony; the SIMR Histology Core, Cindy Maddera, the SIMR Genome Engineering core and the SIMR Sequencing Core for experimental assistance; Sofia Robb for OMA support; the Stowers Café for providing lettuce for feeding the snails. The work was funded by HHMI (ASA), SDB (AA), AAA (AA) and by institutional support from the Stowers Institute for Medical Research (ASA).

## AUTHOR CONTRIBUTIONS

Conceptualization: AA and ASA; Methodology: AA, TJC, KD; Investigation: AA, TJC, MM, KW, JM, MCM; Data Analysis: AA, ER, BP, SAM; Writing – original draft: AA; Writing – review and editing: AA, ER, BP, TJC, MM, KW, KD, JM, MCM, SAM, ASA; Funding acquisition: AA, ASA; Resources: ASA; Supervision: ASA.

## DECLARATION OF INTERESTS

The authors declare no competing interests.

### Materials, data and code availability

Apple snail lines generated in this study will be made available upon request, but we may require a payment and/or a completed materials transfer agreement if there is potential for commercial application.

Original data underlying this manuscript can be accessed from the Stowers Original Data Repository at http://www.stowers.org/research/publications/libpb-2417. Eye regeneration RNA-Seq data have been deposited at GEO with accession number GSE240085. Tissue-specific RNA-Seq dataset was previously deposited as a BioProject with accession number PRJNA473253. All original code has been deposited in GitHub at https://github.com/AliceAccorsi/SnailEyeRegeneration. Any additional information required to reanalyze the data reported in this paper is available from the lead contact upon request.

### Animal husbandry

The *Pomacea canaliculata* colony at the Stowers Institute for Medical Research (Kansas City, USA) was established starting from specimens shipped from the University of Modena and Reggio Emilia (Modena, Italy) where a colony of apple snails was maintained for more than 6 years. The apple snails were maintained at 25-27 °C on a light-dark cycle of 14 and 10 h, respectively. They were fed three times a week with a variety of green leaves and kale or calcium enriched pellets to support shell growth. System water was the result of deionized water (diH_2_O) mixed with a variety of salts (2.15 uM CaCl_2_, 0.95 uM MgSO_4_, 1.88 uM NaHCO_3_ and 0.02% P-Remineraliz^TM^ (Brightwell® Aquatics, USA)) to reach pH = 7.5, osmolarity = 1100 uS, alkalinity 69 ppm, carbonate hardness 248 ppm, 71.53 mg/l Calcium, 21.97 mg/l Magnesium, 2.07 mg/l Potassium, 87.32 mg/l Sodium, 77.36 mg/l Sulfate and additional elements in lower concentrations. pH was maintained constant through a Sodium Bicarbonate automatic dosing system. Additionally, daily water changes (10-30% of total water volume) and debris siphoning helped maintaining good water quality.

### Adult dissection

Unless differently specified, we used adult apple snails with shell length (A-P axis) ≥ 27 mm and from 3 to 7 months old. To access their cephalic area, the apple snails were anesthetized through a 10 min incubation in ice and then the operculum was gently pulled with self-clamping forceps. Surgical micro-scissors and tweezers were used to fully remove the eyes, to collect intact or regenerating tissue or to isolate the lens or the retina. Images were acquired with a Zeiss Stemi SV 11 Apo stereomicroscope equipped with a Lumen*era* Infinity3 camera or with a Leica M205 FA stereomicroscope equipped with a Leica DFC310 FX camera. After image acquisition, Fiji software^60^ was used for exporting and processing images.

### Embryo collection

To collect embryos in the first 24 h post fertilization (hpf), clutches were incubated in 15 mg/ml L-cystein pH 7.5 for 3 min. Then, they were moved in Petri dishes while the L-cystein solution was diluted 1:1 with Embryo Media (6 mM KCl, 6.7 mM CaCl_2_, 3.3 MgCl_2_, 1.7 mM HEPES, 33.62 mM NaCl in diH_2_O). We individually opened all the capsules with tweezers and then alternated gentle swirls of the mix with embryo harvesting using a stereomicroscope. The gentle swirls helped the embryos to be released from the perivitelline fluid (PVF). To collect embryos after 24 hpf, the capsules were crushed in Embryo Media with tweezers in dishes. Gentle swirls of the media helped to release the embryos from the PVF. Embryos sink on the bottom of the dish at all stages. Since embryos younger than 4 dpf stick to plastic and glass, 5% FBS was added to Embryo Media.

### Hematoxylin & Eosin (H&E) morphological staining

Immediately after collection, the samples were fixed in 4% paraformaldehyde (PFA) at 4 °C overnight (ON) on a rotator. After fixation, the samples were incubated twice in 1x PBS with 0.5% Triton X-100 (PBSTx 0.5%) for 20 min, in 50% MeOH in diH_2_O for 10 min and in 100% MeOH for 10 min. Finally, the samples were stored in clean 100% MeOH at -20 °C.

To embed the samples in paraffin, we processed them using a Milestone PATHOS Delta Microwave Tissue Processor. The incubations we performed were: 70% EtOH for 4 min, 100% EtOH for 10 min at 65 °C, 100% isopropyl alcohol for 45 min at 68 °C, Paraffin Type 9 (Epredia #8337) at 70 °C for 10, 10, 2, 2 and 2 min, and finally in Paraffin Type 9 at 65 °C for 27.5 min. The samples were stored at room temperature (RT) in cassettes until they were ready to be oriented and embedded in paraffin using a LEICA EG1150H Embedding Center. The samples were sectioned at a 7-8 um thickness onto charged glass slides (Fisher Scientific #22-042-924) using an Automated Rotary Microtome (HistoCore AUTOCUT #149AUTO00C1 and #14051956472) and an Illuminated Tissue Bath set at 42 °C (230 V) (Boekel Scientific #145951-2). Slides were dried in a dry oven at 36 °C for at least 4 h and then stored at RT.

Sections were deparaffinized and stained with ST Infinity H&E staining system kit (Leica #3801698) using a Leica ST5010 Autostainer XL. Slides were baked for 60 min at 60 °C, then they were incubated: three times in 100% xylene over 9 min, three times in 100% EtOH over 3 min, in 80% EtOH for 1 min, in diH_2_O for 1 min, in Hemalast for 30 sec, in Hematoxylin for 2 min, in diH_2_O for 2 min, in Differentiator for 45 sec, in diH_2_O for 1 min, in Bluing for 1 min, in diH_2_O for 1 min, in 80% EtOH for 1 min, in Eosin for 30 sec, three times in 100% EtOH over 9 min and three times in 100% xylene over 3 min. Finally, the coverslips were mounted with Surgipath Micromount (Leica #3801731) using a Leica CV5030 Fully Automated Glass Coverslipper and dried for 15 min at RT.

Images were acquired with a Leica DM6000 B upright microscope equipped with a Leica DFC425 camera or with an Olympus SlideScanner VS140 and a 40x objective. After image acquisition, Fiji software^60^ was used for exporting and processing images and measuring the eye components.

### Immunohistochemistry (H3P/BrdU)

Snails were anesthetized and eyes were fully removed as already mentioned. Snails were anesthetized again 24 h before collecting the regenerating tissue and were injected on the left side of the body with 3 mg of BrdU (Sigma #B5002) diluted in 500 ul of diH_2_O. The regenerating tissue was collected 24 h post injection (hpi) and fixed, washed, dehydrated, embedded in paraffin and sectioned as already mentioned.

Sections were deparaffinized using a Leica ST5010 Autostainer XL by baking the slides for 60 min at 60 °C and by incubating them: three times in 100% xylene over 9 min, three times in 100% EtOH over 3 minutes, in 80% EtOH for 1 min and in diH_2_O for 1 min. Sections were bleached for 1 h and 30 min under direct light in 5% formamide, 1% H_2_O_2_ and 0.5x SSC (1x SSC is 150 mM sodium chloride and 15 mM sodium citrate, pH 7.0 with citric acid) in diH_2_O. After rinsing 3 times in diH_2_O, we performed heat-induced antigen retrieval through an incubation in citrate buffer (0.1 M citric acid and 0.1 M trisodium citrate in diH_2_O, pH 6.0) at 95 °C for 15 min in a BioGenex EZ-Retriever^®^ IR – Antigen Retrieval System. Slides were allowed to cool down for 20 min and then they were incubated in 1x PBS with 0.1% Tween 20 (PBSTw 0.1%) for 30 min. Blocking of nonspecific antibody binding was performed using Background Buster (Innovex #NB306) for 45 min at RT. Slides were washed three times with PBSTw 0.1% over 15 min and then incubated with 1:300 Rabbit anti-H3P (Millipore #05817R clone 63-1C-8) and 1:300 Mouse anti-BrdU (BD Biosciences #347580 clone B44) for 1 h at RT. Slides were washed three times with PBSTw 0.1% over 15 min and then incubated with 1:300 Alexa488 Donkey anti-Rabbit (Biotium #20015), 1:300 Alexa568 Donkey anti-Mouse (Biotium #20105) and 1:500 DAPI (Biolegend #422801) in Background Buster for 90 min at RT. Slides were washed three times with PBSTw 0.1% over 15 min and then the coverslip was mounted using Prolong Gold mounting media (Thermo Fisher Scientific #P36984).

Images were acquired with a Zeiss LSM 700 confocal and images were processed using custom plugins and macros in Fiji software^60^ similar to the procedure in previous reports^61^.

### Electron microscopy - TEM processing

Immediately after collection, samples were fixed in EM fixative (2% glutaraldehyde and 2.5% paraformaldehyde in 50 mM sodium cacodylate buffer, with 1 mM calcium chloride and 1% sucrose, pH 7.35) for 1 h at RT on a rotator. After fixation, the samples were stored in EM fixative at 4 °C.

All the following steps were completed at 4 °C unless otherwise noted and all rinsing steps were 15 min each. Samples were rinsed 3 times with 50 mM sodium cacodylate buffer followed by secondary fixation with buffered 1% osmium tetroxide for 1-2 h at RT. Three more rinses with 50 mM sodium cacodylate buffer and three with water were carried out before *en bloc* staining with 0.5% aqueous uranyl acetate at 4 °C ON. After rinsing three times with water again, samples were dehydrated in a graded series of acetone (25%, 50%, 75%, 90% and 100%) for 10 min each. Cold 100% acetone was used for an additional incubation and moved to RT for 10 min, followed by a final 100% acetone incubation at RT for 10 min. Resin infiltration was carried out at RT with a graded series (1:2, 1:1, 2:1) of SPURR low viscosity resin (EMS #1430 hard formulation) with acetone for 30 min to ON each step depending on sample size, followed by 4 pure resin incubations over 2 days. Samples were embedded in SPURR resin in flat molds (Ted Pella #105) and cured at 60 °C for 48 h.

Once cured, sections were cut between 60 nm and 100 nm with a Leica UC6 or UC7 ultramicrotome and Diatome ultra 45 diamond knife and placed on Formvar^®^/Carbon coated slot grids or on glass slides. Sections were then post-stained at RT with Sato’s triple lead stain for 3 min, 4% uranyl acetate in 70% MeOH for 4 min and Sato’s triple lead stain again for 5 min. Grids were imaged in a Tencnai BioTwin TEM at 80 kV with a Gatan UltraScan 1000 CCD camera, or a Zeiss Merlin SEM using aSTEM4 detector at 30 kV and 141 pA with either SmartSEM (Zeiss) or Atlas 5 (Fibics, inc.) acquisition software. Slides were coated with 4 nm of carbon in a Leica ACE600 and imaged in a Zeiss Merlin SEM with a 4 quadrant BSD detector at 8 kV and 700 pA using Atlas 5 acquisition software.

For vesicle analysis, sections were cut at 60 nm on slides and sections post-stained and coated with carbon as already mentioned. For each eye an overview of one section at 100 nm pixel size and 2 areas of interest at 6 nm pixel size was acquired on a Zeiss Merlin SEM as stated above using Atlas 5 software.

Acquired images were exported and processed using Fiji software^60^. Photic vesicles were identified using the deep learning package DeepFiji^62^ after hand annotating examples.

### Electron microscopy - SEM processing

Immediately after collection, the samples were fixed and stored as already mentioned.

All the following steps were completed at RT and all rinsing steps comprised 2 quick rinses and then 5 rinses over 25 min. Staining and secondary fixation solutions were all 1% aqueous and filtered with a 0.2 um syringe filter. Samples were post-fixed and stained with tannic acid, osmium tetroxide, thiocarbohydrazide (TCH), and osmium tetroxide sequentially, each for 1 h and rinsed in ultrapure water before each incubation. After rinsing again with ultrapure water, samples were dehydrated in a graded series of EtOH (30%, 50%, 70% and 90%) and 3 pure EtOH incubations for 10 min each. Samples were loaded into a 4 or 12 sample holder (Tousimis #8763 or #8762) for critical point drying in a Tousimis Samdri-795. After drying, samples were mounted on aluminum stubs with carbon stickers, coated with 4 nm gold palladium in a Leica ACE600, and imaged in a Zeiss Merlin SEM at 8 kV and 150 pA with SE detector and a working distance of 8 mm, using SmartSEM or Atlas 5 acquisition software.

Acquired images were exported and processed using Fiji software^60^.

### Bulk RNA-Sequencing (RNA-Seq)

Adult tissues and organs were collected and total RNA from each sample was extracted using Trizol^®^ Reagent (Ambion #15596026) and following the protocol from the producer. The purified RNA quantity and quality was determined by Qubit^®^ 2.0 Fluorometer (Invitrogen) and Agilent 2100 Bioanalyzer (Agilent Technologies), respectively.

mRNA-Seq libraries were generated from high-quality total RNA as assessed using the Bioanalyzer (Agilent) or the LabChip GX (Caliper Life Sciences). Libraries were made according to the manufacturer’s directions for the TruSeq Stranded mRNA LT Sample Prep Kit – sets A and B (Illumina #RS-122-2101 and #RS-122-2102) or TruSeq Stranded mRNA Library Prep kit (48 Samples) (Illumina #20020594) or TruSeq RNA Single Indexes Sets A and B (Illumina #20020492 and #20020493). Resulting short fragment libraries were checked for quality and quantity using the Bioanalyzer (Agilent) or the LabChip GX (Caliper Life Sciences) and the Qubit Fluorometer (Life Technologies). Libraries were pooled, re-quantified and sequenced utilizing RTA and instrument software versions current at the time of processing.

In more details, mRNA-Seq libraries for retinas were generated from 250 ng of high-quality total RNA and libraries were sequenced as 50 bp single reads on 1 lane of an Illumina HiSeq 2500 flow cell.

mRNA-Seq libraries for cephalic tentacles were generated from 150 ng of high-quality total RNA and libraries were sequenced as 100 bp paired reads on 4 Rapid Run flow cells using the Illumina HiSeq 2500 instrument.

Samples of intact eyes and regenerating tissue after full eye bulb amputation were collected pooling together tissue from multiple animals. mRNA-Seq libraries for these samples were generated from 100 ng of high-quality total RNA and libraries were sequenced as 50 bp single reads on a P3 flow cell using the Illumina NextSeq 2000 instrument.

mRNA-Seq libraries for all the other samples were generated from 200 ng of high-quality total RNA and libraries were sequenced as 100 bp paired reads on 6 Rapid Run flow cells using the Illumina HiSeq 2500 instrument as well as 150 bp paired reads on 2 Illumina MiSeq flow cells.

### RNA-Seq analyses

RNA-Seq reads for all experiments were aligned to the REFSEQ genome and gene models for *P. canaliculata* (GCF_003073045.1) using the STAR alignment program (v2.7.3a)^63^. TPMs were calculated using RSEM (v1.3.0)^64^.

The relationship between RNA-Seq samples was assessed with MDS and Spearman correlation on normalized counts. MDS plot was generated with all genes and the Spearman correlation plot with the top 1000 genes with highest mean across time points. Differential Expression Analysis (DEA) was performed using the R package edgeR^65–67^. DEA was performed using 1) the intact eye as a reference, 2) the 1 dpa eye as a reference and 3) in a pairwise manner with adjacent timepoints using the earlier time point as the reference for the later one. Differentially expressed genes were defined by setting an FDR cutoff of 1e-5 and a logFC of 0. Additional data manipulation was performed with the tidyverse R package collection^68^.

To perform Gene Ontology (GO) Enrichment^69^, it was necessary to annotate the REFSEQ genes with GO terms. Terms used were derived by combining GO terms from Interproscan (5.42-78.0) results and OMA (v2.4.2) mappings^70–72^. Annotations were put into an “org” file using the AnnotationForge R library^73^.

GO Enrichment analysis was performed with R package clusterProfiler^74,75^. Figures were made using R libraries GOsemsem, cowplot and enrichplot^76–78^. The *p* value cutoff for term enrichment was 0.05 and a *q* value of 0.01. GO terms were explored using GO-Figure!’s semantic similarity scatterplots^79^ and manually selected for display.

### Ortholog analysis

Domain prediction and domain images for genes annotated as *pax6* in *Homo sapiens*, *Drosophila melanogaste*r and *P. canaliculata* were created using SMART^80^. Structures for *pax6* genes were generated with ColabFold (v1.5.2)^81,82^ and visualized with ChimeraX^83^. Multiple sequence alignment was made with MUSCLE^84,85^. OMA (v2.5.0) was used to identify ortholog pairs for comparison of *P. canaliculata* genes to *H. sapiens* and *D. melanogaste*r^70^.

### *Ex ovo* embryo culture

To prepare the perivitelline fluid extract (ePVF) for culturing the embryos *ex ovo*, clutches (1 or 2 dpf) were incubated in 10% bleach for 3 min and then rinsed three times with diH_2_O. The clutches were moved on a dry Petri dish, individually crushed with tweezers, moved in a 2 ml tube and centrifuged at 4 °C for 40 min at 21,000 x*g*. The supernatant was moved in a 1.5 ml tube and stored at -20 °C. The PVF extract can be frozen and thawed multiple times. Because of the nutritious nature of the PVF, autoclaved plastics and clean surfaces help to reduce contaminations. Each ePVF aliquot, coming from an individual clutch, was tested for contaminations and for extract quality using wild-type embryos. For this test, clutches (1 or 2 dpf) were incubated in 10% bleach for 3 min and then rinsed three times with diH_2_O. Embryos were collected as already mentioned and then washed three times in 5% FBS in Embryo Media. It is important to keep clutches separate and process them individually.

To set up an embryo culture, the lid of a 30-mm Petri dish was placed facing up inside a 60-mm Petri dish. Drops of ePVF extract were placed inside the 30-mm Petri dish lid and 3-4 embryos were pipetted inside the ePVF extract drop. If the drops are too small or if too much Embryo Media is released in the drops together with the embryos, the culture might not work. Finally, the lid was filled with about 3.0-3.5 ml of paraffin oil (MilliporeSigma #PX0047-1) to cover all the drops. The *ex ovo* cultures were stored inside 60-mm Petri dishes at RT in the dark and they were checked regularly for contamination, embryo growth or formation of bubbles that could cause them to dry out.

### Zygote micro-injections

Embryos were harvested as soon as possible as already mentioned and then split into two Petri dishes in 5% FBS in Embryo Media. One dish was placed at RT and one at 4 °C to slow down the embryo development. For the micro-injection procedure, 15-20 zygotes were taken from the Petri dish at RT and placed on a glass depression slide. The Petri dish at RT was then moved at 4 °C, while the other one was moved from 4 °C to RT for the next round of micro-injection. This alternating pattern was continued for subsequent rounds until all the embryos were micro-injected. The glass slide with zygotes was placed on a cooling stage at 12 °C located on a Nikon Eclipse Ti2 inverted microscope equipped with Eppendorf TransferMan 4r micromanipulators. The cooling stage increased the rigidity of the embryos. Micro-injection was performed using a 20x objective and DIC optics. Embryos were held in place with a standard holding pipette made from borosilicate glass (Harvard Apparatus #GC100-15). Holding pipettes were pulled with a Sutter Horizontal needle puller P-87, flame polished with a Narishige microforge, and connected to an Eppendorf CellTram 4r Air pneumatic syringe. The holding pipette kept the embryos still and facilitated pulling the needle out. Embryo injections were performed using glass needles made from borosilicate glass capillaries containing an internal filament (Harvard Apparatus #GC100TF-10) fashioned with a Sutter Horizontal needle puller P-87 and backfilled by capillary pressure. The tip of the glass needle was gently opened using the edge of the holding pipette. Too small/sharp of a tip will result in immediate lysis of the embryos while too dull of a tip will not allow the needle to penetrate the cell membrane.

Micro-injection pressures were controlled with an Eppendorf FemtoJet 4i (compensation pressure 20-30 hPa, injection pressure 90-100 hPa). Additionally, the WPI MICRO-ePORE pinpoint cell penetrator electroporation device was incorporated into the micro-injection procedure (frequency 520 Hz, amplitude 0.480 V). The MICRO-ePORE reduced the mechanical stress of the micro-injection. Following micro-injection, the embryos were moved from the slide to a new Petri dish with 5% FBS in Embryo Media and stored at RT.

To test for micro-injection efficiency, we injected 2 mg/ml 3 kDa Dextran Texas Red (Thermo Fisher Scientific #D3329) and 0.05% Phenol Red (Sigma #P0290) in diH_2_O.

To synthesize exogenous mRNA, we amplified the constructs of interest through PCR with Phusion High-Fidelity DNA Polymerase (NEB #M0530), we purified the PCR product, we performed *in vitro* transcription using the mMESSAGE mMACHINE High Yield Capped RNA Transcription Kit (Invitrogen #AM1344 or #AM1348) following the manufacturer’s protocols, we used the Poly(A) Tailing Kit (Invitrogen #AM1350) and we purified the synthesized mRNA with the MEGAclear Transcription Clean-Up Kit (Invitrogen # AM1908). After quantifying the mRNA using the Qubit^®^ 2.0 Fluorometer (Invitrogen), we micro-injected the embryos with 75 ng/ul exogenous mRNA and 0.05% Phenol Red in diH_2_O.

The dividing embryos were cultured *ex ovo* using the ePVF aliquots that passed the contamination and quality test. After 11-13 days in *ex ovo* culture, the embryos were moved in a Petri dish in Hatchling Media (1 mg/ml galactose and 1 mM CaCl_2_ in Embryo Media). After 3-5 days the snails were moved in Juvenile Media (1 mM CaCl_2_ in Embryo Media) and fed with small pieces of salad and after 7-10 days they were moved in system water.

### Fluorescent *in situ* hybridization (HCR)

After collecting the tissue of interest as already mentioned, we fixed the samples in 4% PFA with 0.1% Tween 20 at 4 °C ON on a rotator. After fixation, the samples were incubated twice in PBSTx 0.5% over 20 min, then, in pre-heated reduction solution (1% NP40 or IGEPAL, 0.5% SDS in 1x PBS; 50 mM DTT was added only before use) at 37 °C for 15 min, twice in PBSTx 0.5% over 20 min, in 50% MeOH in diH_2_O for 10 min and in 100% MeOH for 10 min. Finally, the samples were stored in clean 100% MeOH at -20 °C.

To re-hydrate and bleach the samples, we incubated them in 50% MeOH in diH_2_O for 10 min, in PBSTw 0.5% for 10 min, twice in 0.5% Tween 20 in diH_2_O over 10 min, in 80% acetone in diH_2_O for 30 min at -20 °C, four times in PBSTx 2% for 30 min and finally in bleaching solution (6% H_2_O_2_ in PBSTx 0.5%) under direct light. To store the samples, we repeated the washes and de-hydration steps already performed after the PFA fixation.

Fluorescent *in situ* hybridization chain reaction (HCR) v3^86^ was performed based upon Molecular Instruments’ RNA-FISH generic samples in solution protocol. If the samples are too small to settle by gravity, we centrifuged at 500 x*g* for 10 sec. Samples were processed in 1.7 ml or 2 ml tubes that were positioned horizontally and allowed to rock gently during all incubations and washes.

HCR was performed on *pax6*-B5 (LOC112559942), *rhodopsin*-B4 (LOC112576458) and *nec2*-B2 (LOC112570448) (Molecular Instruments) and the amplifiers used were B2-488, B4-546 and B5-647 (Molecular Instruments).

Immediately prior to imaging, samples were cleared in OptiPrep (Iodixanol) (Sigma #D1556) for 30-60 min at RT and then mounted in ProLong Glass Mountant (Thermo Fisher Scientific #P36980) between two coverslips separated by vacuum grease. Images were acquired with a Nikon CSU-W1 Ti2-E Spinning Disk confocal microscope. After image acquisition, Fiji software^60^ was used for exporting and processing images.

### Stable mutant lines through CRISPR-Cas9

Genomic locations of interest were evaluated for potential guideRNA (gRNA) target sites using CCTop^87^. The resulting sites were assessed using the predicted on-target efficiency score and the off-target potential^88^. The gRNA had to rank medium or high from CCTOP and target a location downstream of the start site and inside of an exon. The sequences were ordered as Alt-R CRISPR-Cas9 crRNA from Integrated DNA Technologies (IDT). Five ul of 500 ng/ul crRNA was hybridized with 10 ul of 500 ng/ul universal tracrRNA (IDT) at 95 °C for 5 min and then cooled to RT to form a full-length gRNA. The ribonucleoprotein (RNP) complex was prepared by mixing 1 ul of 500 ng/ul hybridized cr/tracrRNA with 0.5 ul of 1 ug/ul Cas9 protein HiFi V3 (IDT), incubating at RT for 20 min, cooling on ice for 20 min and then adding 43.5 ul of diH_2_O and 5 ul of 0.5% Phenol Red (Sigma #P0290). The injection mix was centrifuged at 20,000 x*g* for 5 min at 4 °C and then the top 35 ul were recovered in a new tube. The final injection mix composition was 10 ng/ul hybridized cr/tracrRNA, 10 ng/ul Cas9 protein HiFi V3 and 0.05% Phenol Red in diH_2_O stored on ice until use.

Embryos were micro-injected, *ex ovo* cultured and grew to adults as already mentioned. Some F0s were genotyped to evaluate the gRNA activity and efficiency of this technology on apple snails. Other F0s were in-crossed with each other or out-crossed with wild-type snails. F1s from each cross were genotyped and then selected F1s were in-crossed to obtain F2s. Finally, to maintain the lines, F1s were out-crossed with wild-type snails and the offspring was genotyped and selected based on presence of indels.

To genotype apple snails, whole F0 or F1 embryos or tips of tentacles were lysed in 50 ul of QuickExtract^TM^ DNA Extraction Solution (Epicentre #QE09050) and processed as suggested by the manufacturer (65 °C for 6 min and 98 °C for 2 min). PCR was performed (98 °C for 30 sec; 30 times 98 °C for 10 sec, annealing temp. for 10 sec, 72 °C for 10 sec; 72 °C for 2 min) to amplify the specific genomic location, followed by a second round of amplification to incorporate sample-specific dual barcodes. Each PCR reaction was set up with 5 ul NEB Next enzyme (New England BioLabs), 1 ul of lysed tissue and 0.5 uM of each primer. All amplicons were pooled and size-selected using ProNex Size-Selective Purification System (Promega #NG2001). The quantity and quality of cleaned pools were determined by the Qubit^®^ 2.0 Fluorometer (Invitrogen) and Agilent 2100 Bioanalyzer (Agilent Technologies), respectively. Purified pools were run on an Illumina MiSeq 250 flow cell. The resulting sequence data was demultiplexed, and read pairs were joined. On-target indel frequency and expected mutations were analyzed using CRIS.py^89^.

### Embryo phenotype analyses

To evaluate phenotypes, embryos were harvested at the desired stage as already mentioned. They were mounted on a slide using 2% methylcellulose in Embryo Media and imaged with a Leica M205 FA stereomicroscope equipped with a Leica DFC310 FX camera. After image acquisition, Fiji software^60^ was used for exporting and processing images.

Embryos were also fixed in 4% PFA at 4 °C ON, washed four times in PBSTx 2% over 2 h, washed twice in PBSTx 0.5% over 10 min and incubated in 1:400 AlexaFluor488-Phalloidin (Invitrogen #A12379) in PBSTx 0.5% for 2 h at RT. We washed the samples three times in PBSTx 0.5% over 15 min, incubated in 1:1000 DAPI in PBSTx 0.5% and rinsed once in 1x PBS. After washing the embryos twice in 1x PBS, we mounted them on glass bottom Petri dishes using 1% low melting point agarose in Embryo Media and acquired confocal images using the Nikon CSU-W1 Ti2-E Spinning Disk confocal microscope. After image acquisition, Fiji software^60^ was used for exporting and processing images.

### Statistical analysis

All statistical tests were performed using GraphPad Prism 9 (version 9.4.1). Normal distribution was assessed using Kolmogorov-Smirnov test. Statistical significance was calculated using either one-way ANOVA with Tukey’s post hoc multiple comparisons test or non-parametric Kruskal-Wallis ANOVA with Dunn’s post hoc multiple comparisons test. *p* values smaller than 0.05 were considered statistically significant. Plots were made in GraphPad Prism 9.

---

**Figure S1.**
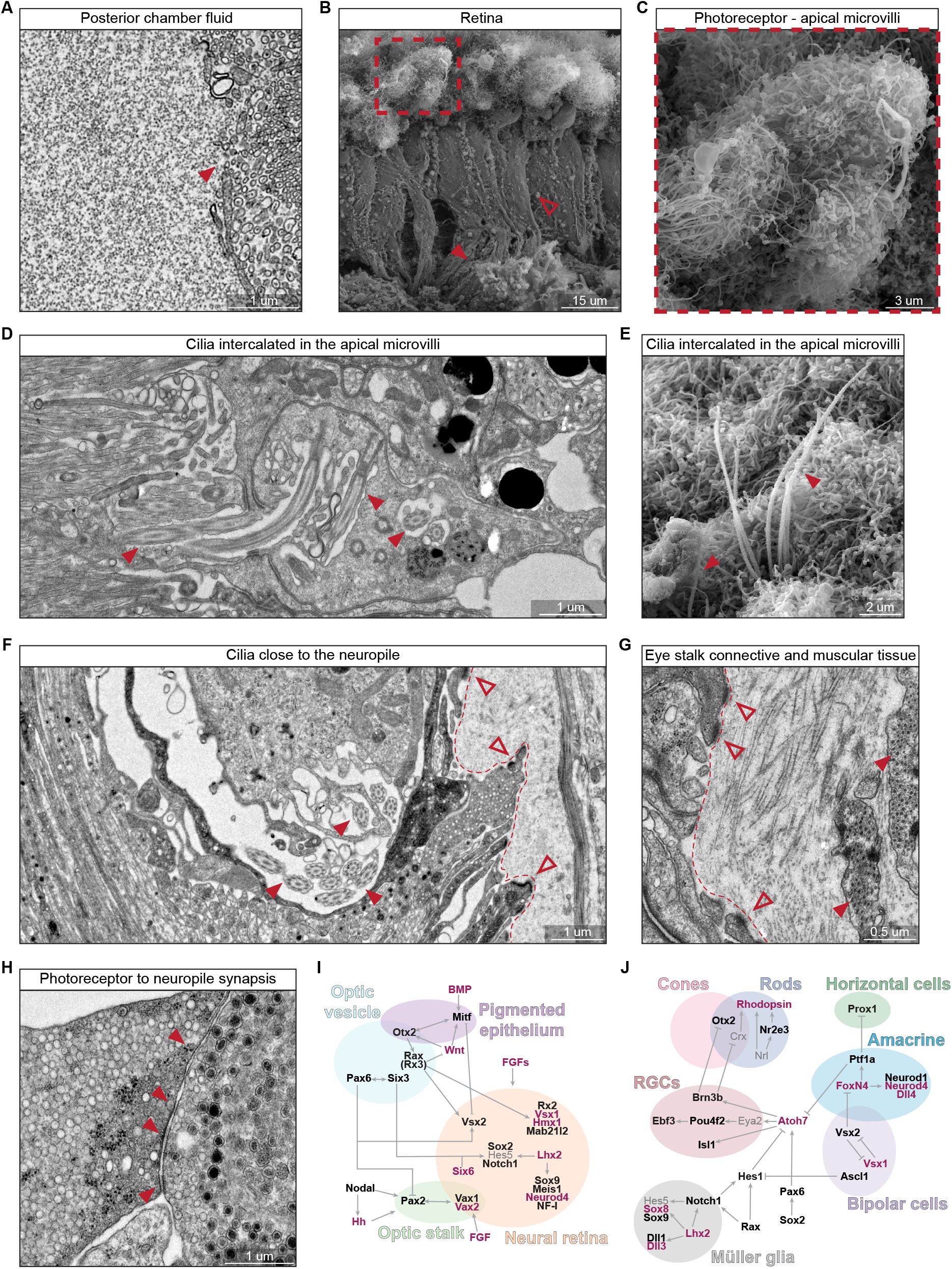
*P. canaliculata* retina has ciliated cells, related to Figure 1. **(A)** TEM image of the posterior chamber fluid (on the left) and of the most apical portion of the photoreceptors microvilli (on the right, full arrowhead). **(B)** SEM image of the retina, constituted by elongated photoreceptors (empty arrowhead), neuropile (on the bottom, full arrowhead) and microvilli (on the top, dashed square). **(C)** Zoomed in SEM image of the photoreceptor microvilli, showing an apical flat and wide expansion of the membrane. **(D)** TEM and **(E)** SEM image of cilia (full arrowhead) intercalated in the photoreceptor microvilli. **(F)** TEM image of multiple cilia (full arrowhead) localized closer to the neuropile and the external edge of the retina (dashed line) defined by retinal cells adhesions (empty arrowheads) with ECM. **(G)** TEM image of the connective and muscular tissue surrounding the eye bulb. The retina cells (on the left) create adhesions (empty arrowheads) with the ECM (central portion of the image). The eye stalk is constituted mainly by ECM and muscle cells (full arrowhead). **(H)** TEM image of a synapsis (full arrowhead) between the basal part of a photoreceptor with photic vesicles (on the left) and the neuropile full of heterogeneous vesicles (on the right). Schematic representation of vertebrate Gene Regulatory Network (GRN) during **(I)** optic cup patterning and **(J)** retinal differentiation (modified from Buono and Martinez-Morales^1^). The black font marks genes that have reciprocal best BLAST hits in *P. canaliculata* genome, the purple font marks genes that belong to a gene family whose members have BLAST hits in *P. canaliculata* genome, the gray font marks genes that have not been identified in *P. canaliculata* genome (see Table S2).

**Figure S2.**
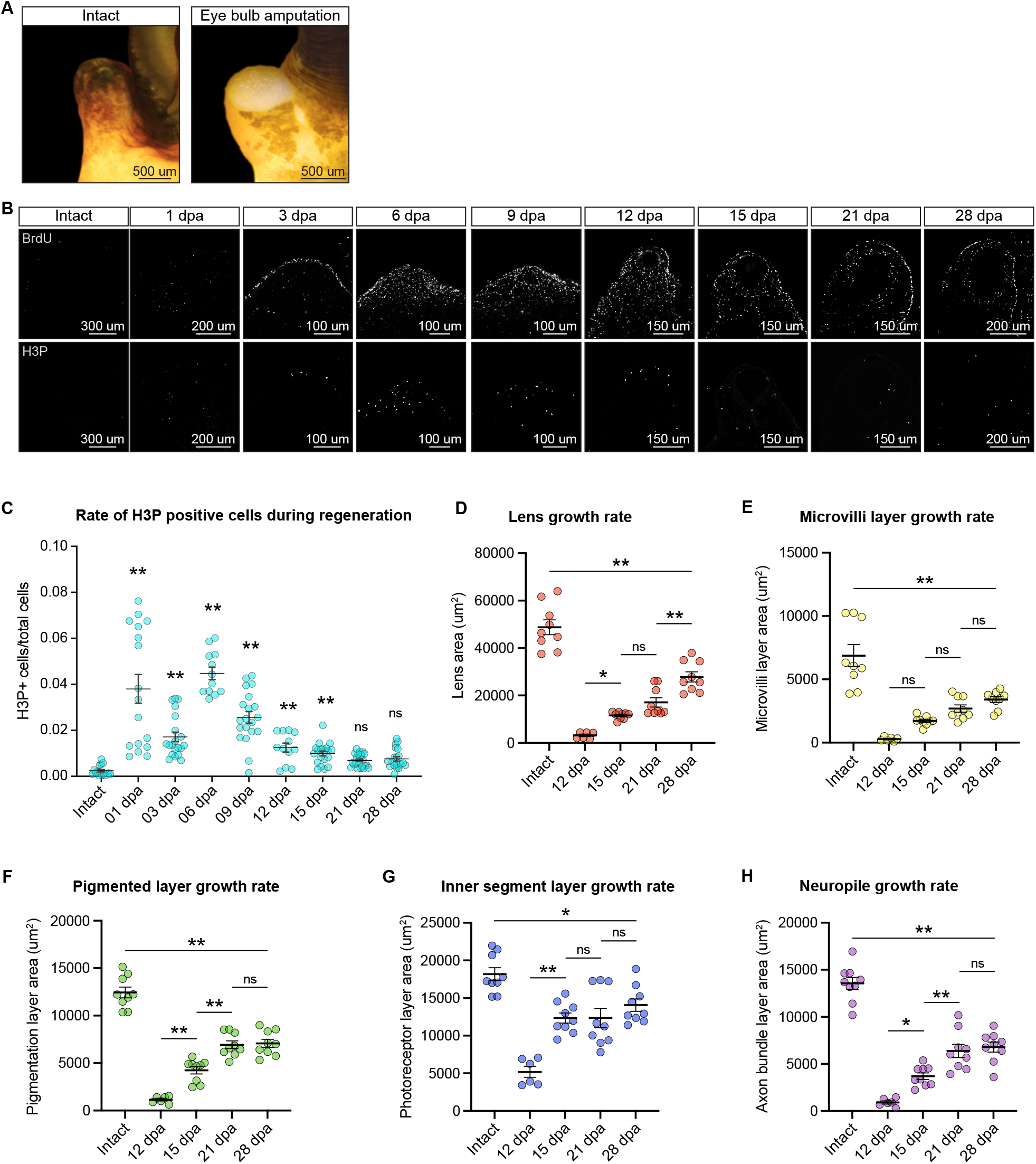
*P. canaliculata* eye components have different growth rate, related to Figure 2. **(A)** Intact eye stalk and open wound immediately after complete amputation of the whole eye bulb. **(B)** Longitudinal sections of the eye stalk during eye regeneration time course stained with BrdU (top row) and anti-H3P (bottom row), showed as individual channels. **(C)** Quantification of H3P positive cells in the regenerating tissue area during the eye regeneration time course. n = 3 to 6 snails/time point, 4 sections each. **(D)** Quantification of the lens area during the eye regeneration time course. n = 3 snails/time point, 3 sections each. **(E)** Quantification of the microvilli layer area during the eye regeneration time course. n = 3 snails/time point, 3 sections each. **(F)** Quantification of the pigmented layer area during the eye regeneration time course. n = 3 snails/time point, 3 sections each. **(G)** Quantification of the inner segment layer area during the eye regeneration time course. n = 3 snails/time point, 3 sections each. **(H)** Quantification of the neuropile area during the eye regeneration time course. Data are represented as mean ± SEM. n = 3 snails/time point, 3 sections each. * = *p* value < 0.05, ** = *p* value < 0.01, ns = non-significant

**Figure S3.**
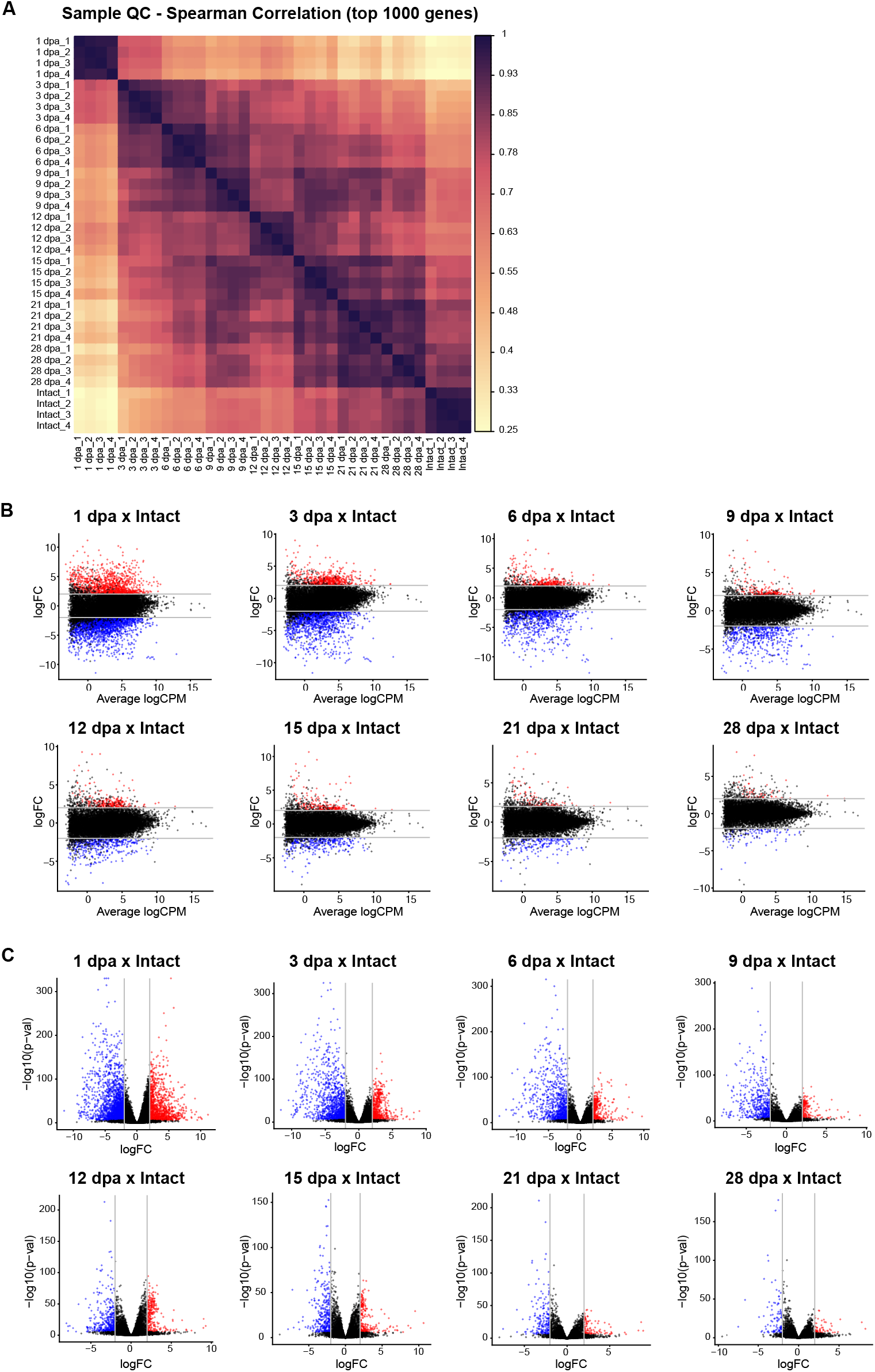
Eye stalk transcriptome changes during *P. canaliculata* camera-type eye regeneration, related to Figure 3. **(A)** Spearman correlation of the top 1000 genes with highest mean expression across all eye regeneration time points. **(B)** MA plots and **(C)** Volcano plots from all regeneration time points compared to Intact eye. Significantly up-regulated genes are in red and significantly down-regulated genes are in blue (FDR <= 1e-5). Gray lines are placed at LogFC threshold +2 and -2.

**Figure S4.**
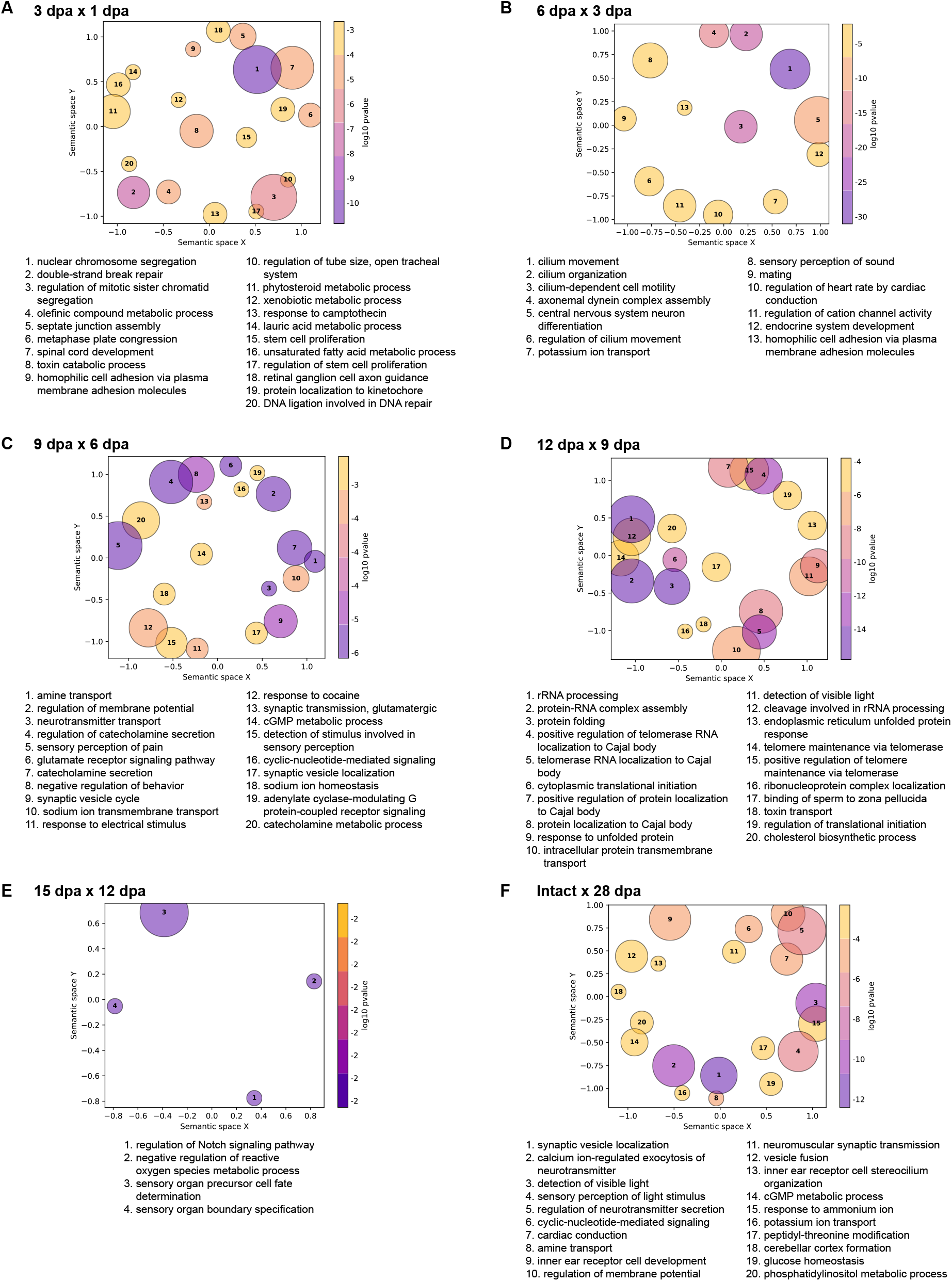
GO enrichment analysis highlights changes during the eye regeneration time course, related to Figure 3. **(A-F)** Summary visualizations of GO term lists obtained from running the GO enrichment analysis on the up-regulated genes from comparing each time point with the previous one. This visualization was generated with the tool GO-Figure!^2^, which simplifies GO term lists by grouping together terms based on information content and semantic similarity. Groups are then plotted in two-dimensional semantic space and labeled with a representative term. Circle size corresponds to the number of terms comprising a given group and color to log10 *p* values of the representative term. Used similarity cutoff is 0.4 and a maximum amount of 20 groups per comparison is shown.

**Figure S5.**
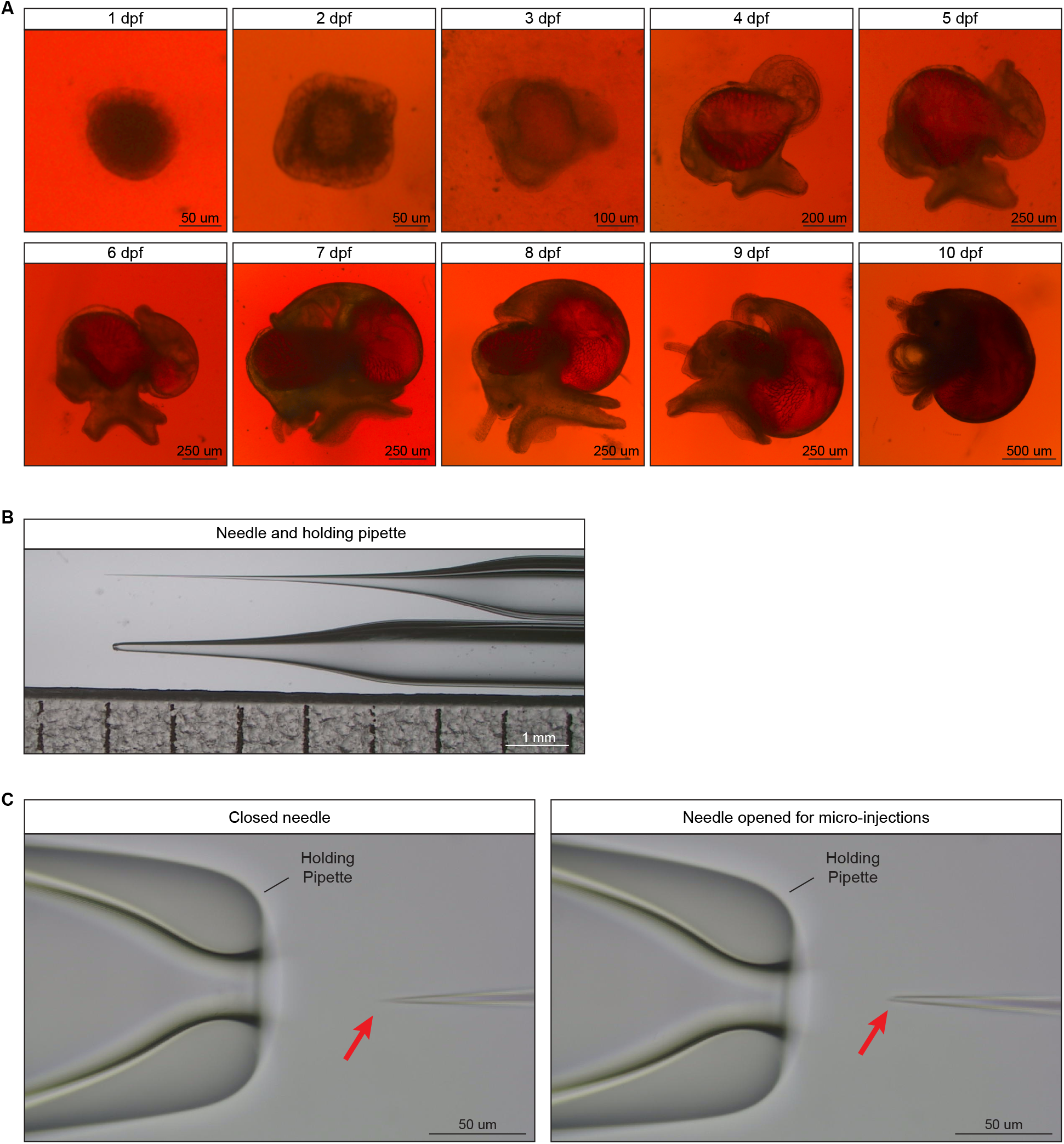
*P. canaliculata* embryos develop *ex ovo*, related to Figure 4. **(A)** Time course of embryos cultured at 0 dpf and developing in the *ex ovo* culture until they are ready to hatch. **(B)** Image of the needle and the holding pipette used for *P. canaliculata* zygote micro-injection. **(C)** Image of the needle and the holding pipette before and after breaking the tip of the needle (red arrow).

**Figure S6.**
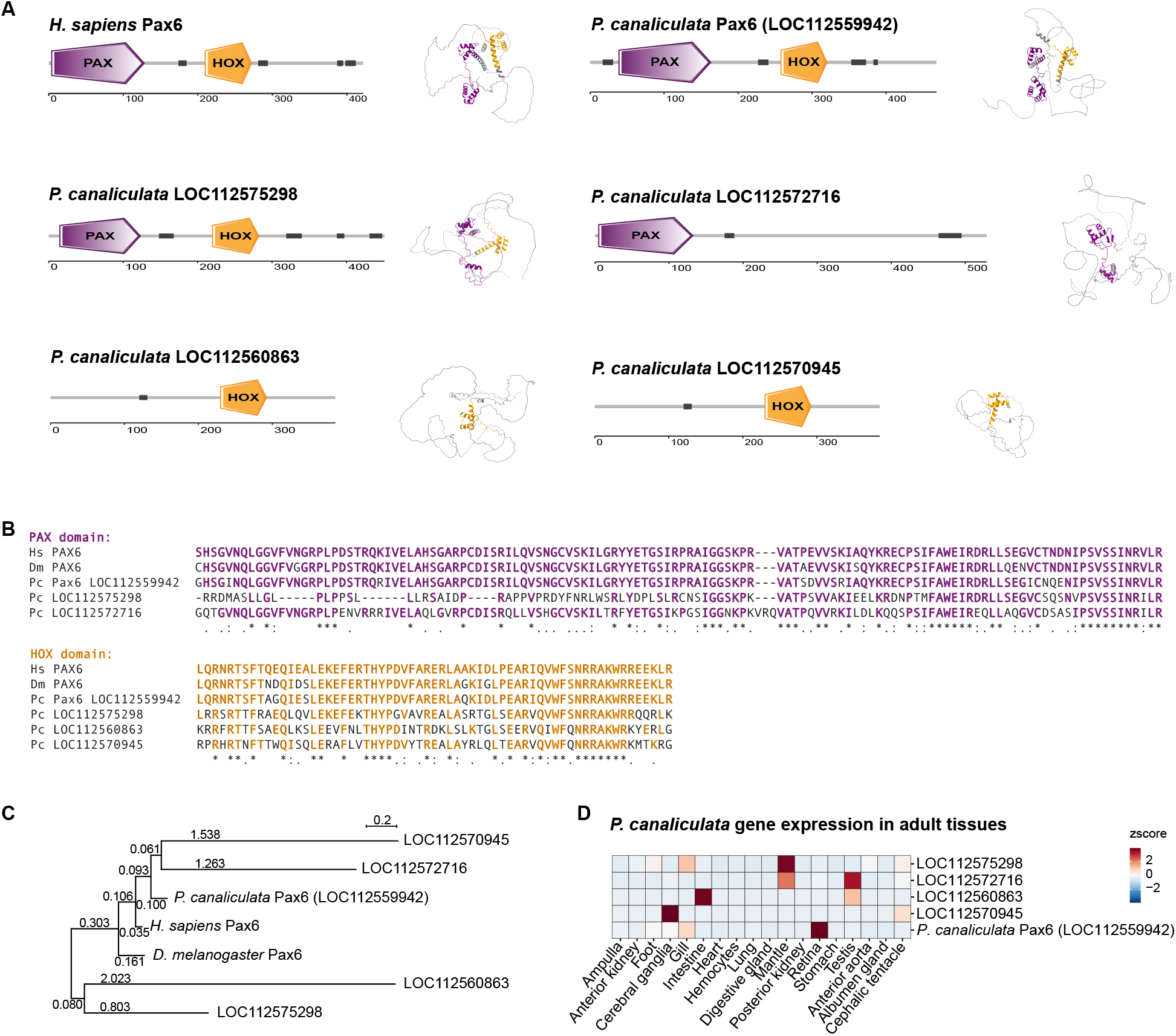
*P. canaliculata* has a gene potentially ortholog of the human and fly *pax6*, related to Figure 5. **(A)** Domain and 3D-folding predictions for human and *P. canaliculata pax6* genes. Purple = PAX domain, orange = HOX domain, dark gray = disordered regions (see Table S4). **(B)** Multi-sequence alignment between human *pax6*, *D. melanogaster pax6* and the 5 genes in *P. canaliculata* annotated as *pax6-like.* Amino acids are bold and colored in purple (PAX domain) and orange (HOX domain) when they are identical to the human amino acid in that specific position. * = sequences have the same residue (fully conserved); : = sequences have residues with strongly similar properties; . = sequences have residues with weakly similar properties. **(C)** Phylogenetic representation of the relationship between human *pax6*, *D. melanogaster pax6* and the 5 genes in *P. canaliculata* annotated as *pax6-like*. The numbers on the branches represent branch lengths. **(D)** Expression levels of the 5 genes in *P. canaliculata* annotated as *pax6-like* in isolated adult tissues, expressed as *z* score.

**Figure S7.**
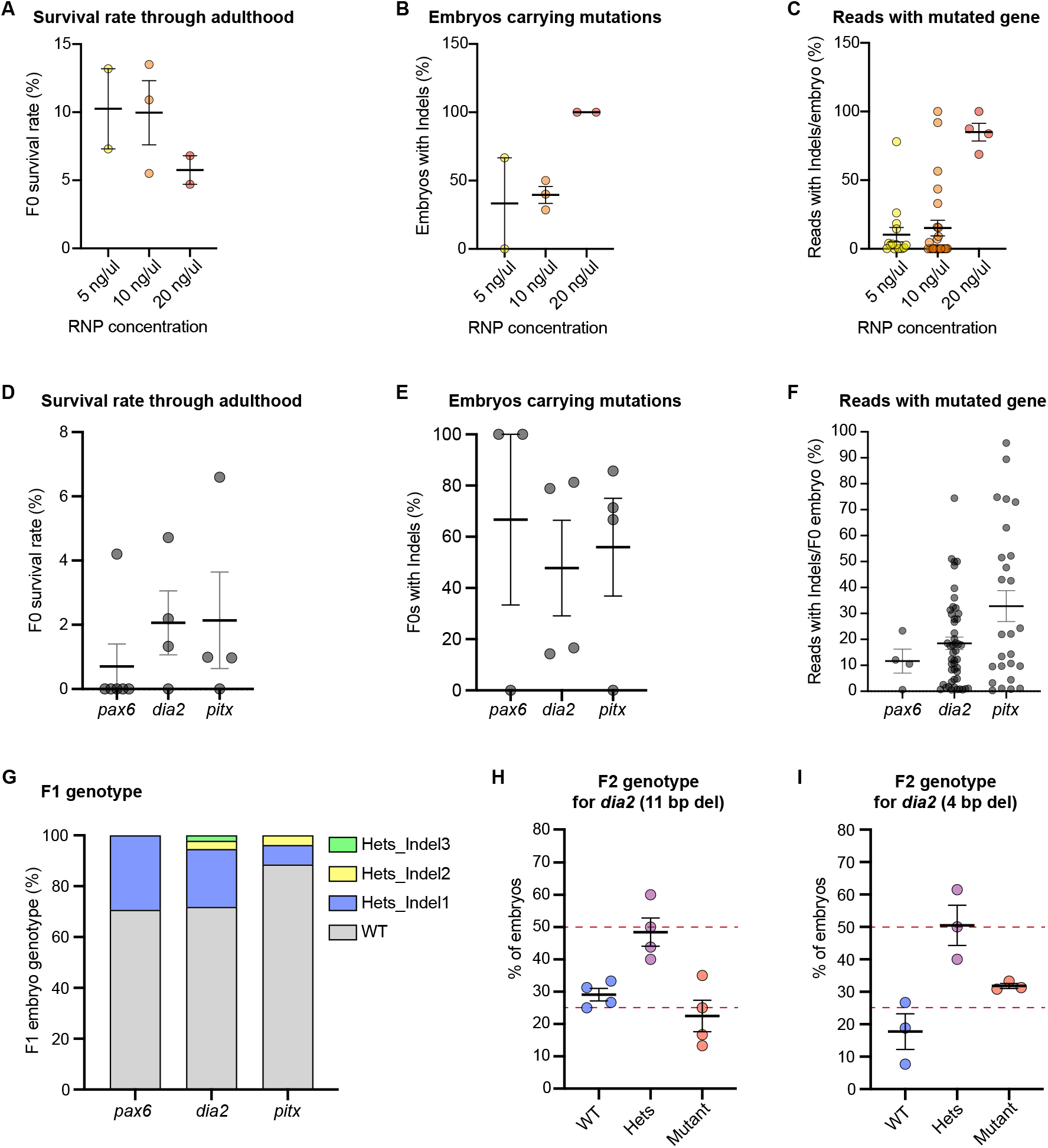
CRISPR-based mutagenesis can now be applied to *P. canaliculata*, related to Figure 6. **(A)** Embryonic survival rate after micro-injection of three different concentrations of RNP (gRNA-Cas9). **(B)** Percentage of embryos with > 10% of the total reads with Indels after micro-injection of three different concentrations of RNP (gRNA-Cas9). **(C)** Percentage of reads with Indels per embryo after micro-injection of three different concentrations of RNP (gRNA-Cas9). **(D)** Embryonic survival rate, **(E)** percentage of embryos with > 10% of the total reads with Indels and **(F)** percentage of reads with Indels per embryo after micro-injection of three different RNPs (gRNA-Cas9) targeting *pax6* (LOC112559942), *dia2* (LOC112560687) and *pitx* (LOC112564278). Each dot represents one injection sessions in (A), (B), (D) and (E) and one embryo in (C) and (F). **(G)** Distribution of F1 embryo genotypes obtained from F0s injected with gRNA targeting *pax6*, *dia2* and *pitx*, respectively. Indel1, Indel2 and Indel3 represent different Indels in term of location and length in each line. We established two mutant lines for *dia2*, one with an 11 bp deletion and one with a 4 bp deletion (see Table S5). The distribution of F2 embryo genotypes obtained for **(H)** the first *dia2* mutant line (11 bp deletion) and for **(I)** the second *dia2* mutant line (4 bp deletion) confirms the expected mendelian frequency of the mutation. n = 4 and 3 clutches, respectively. Data are represented as mean ± SEM.

## SUPPLEMENTAL INFORMATION

**Table S1, related to Figure 1**

Terms obtained from the GO enrichment analysis of *P. canaliculata* (*Pc*) genes bioinformatically defined as orthologs of human (*Hs*) or fly (*Dm*) genes annotated with GO terms related to eye development and function (adjusted *p* value cutoff of 1e-5).

**Table S2, related to Figure S1**

Genes belonging to the vertebrate GRN for optic cup patterning and retinal differentiation^28^, their reciprocal best BLAST hits in *P. canaliculata* genome and their expression during eye regeneration (expressed in TPMs).

**Table S3, related to Figure 3**

Terms obtained from the GO enrichment analyses run on the up-regulated genes obtained from comparing each time point to 1 dpa (first tab) and each time point with the previous one (second tab) (*p* value <= 0.01, q value <= 0.05).

**Table S4, related to Figure S6**

Size, domains and domain conservation of human *pax6*, *D. melanogaster eyeless* and *P. canaliculata* genes annotated as *pax6-like*.

**Table S5, related to Figure 6**

Developed mutant lines through CRISPR/Cas9 technology, genes targeted, gRNA sequences, primers to test for indels and length and position of indels.

**Video S1. Adult *P. canaliculata* shows behavioral phenotype when *pax6* is mutated, related to Figure 6**

Adult *P. canaliculata* free in the tank recorded at 30 frames per second (fps) for about 30 min. Wild-type intact snails, *pax6*^-/-^ animals and wild-type snails at 1 dpa after eye amputation are shown. Videos are played at speed 64x.

## Notes

### Competing Interest Statement

The authors have declared no competing interest.

